# Nanoscopical analysis reveals an orderly arrangement of the presynaptic scaffold protein Bassoon at the Golgi-apparatus

**DOI:** 10.1101/2021.06.28.450125

**Authors:** Tina Ghelani, Carolina Montenegro-Venegas, Anna Fejtova, Thomas Dresbach

## Abstract

Bassoon is a large, 440 kDa, scaffold protein of the specialized sites mediating neurotransmitter release from presynaptic nerve terminals, called active zones. At active zones of the brain, Bassoon is arranged with its C-terminus facing the plasma membrane. In young, cultured neurons Bassoon is associated with the Golgi-apparatus, where it may assemble active zone precursors structures, but whether or not Bassoon is an extended protein at the Golgi-apparatus and whether it is arranged in an orderly fashion is unknown. Understanding the topology of this large scaffold protein is important for models of active zone biogenesis.

Here, we generated recombinant Bassoon constructs for expression in neurons, with tags positioned to allow for the specific detection of N- and C-terminal regions of Bassoon by single-domain antibodies, called nanobodies. Using stimulated emission depletion (STED) nanoscopy in cultured hippocampal neurons, we confirmed that recombinant Bassoon is oriented with its C-terminus towards the active zone plasma membrane at synapses. Focusing on the soma, we found that an intramolecular tag located immediately downstream of amino acid 97 of Bassoon, showed prominent colocalization with markers of the trans Golgi network, including TGN38 and syntaxin-6. In contrast, tags located immediately downstream of the C-terminal amino acid 3938 of Bassoon showed significantly less colocalization with these Golgi-markers. The intramolecular N-terminal tag was located between 48 and 69 nm away from TGN38, while C-terminal tags were located between 100 and 115 nm away from TGN38. Sequences within the first 95 amino acids of Bassoon, but not its N-myristoylation sequence, were required for this arrangement.

Our results indicate that at the Golgi-apparatus Bassoon is oriented with its N-terminus towards and its C-terminus away from the trans Golgi network membrane. Moreover, they suggest that Bassoon is an extended molecule at the trans Golgi network with the distance between amino acids 97 and 3938 estimated to be between 46 and 52 nm. Our data are consistent with a model, in which the N-terminus of Bassoon binds to the membranes of the trans-Golgi network, while the C-terminus associates with active zone components, thus reflecting the topographic arrangement characteristic of synapses also at the Golgi-apparatus.

## 2. Introduction

Scaffold proteins recruit and anchor molecules to subcellular sites. Due to their multi-domain, modular structure, they bind and regulate multiple proteins to coordinate biochemical reactions in space and time. Employing scaffold proteins is a fundamental principle of cell function, operating during protein folding, receptor and signaling molecule clustering, and at cell-cell-junctions (Good et al., 2011).

Synapses are asymmetric cell-cell junctions assembled and regulated by scaffold molecules. On the presynaptic side, a set of synaptic scaffold proteins confines the docking of synaptic vesicles and the exocytotic release of neurotransmitter from these vesicles to specialized sites of the axonal plasma membrane, called active zones. Several families of scaffold proteins operate at active zones, including RIMs, RIM binding proteins, Munc13s, α-liprins and ELKS/CAST/ERC proteins, as well as the particularly large scaffold proteins Bassoon and Piccolo (Sudhof, 2012; Gundelfinger et al., 2016). One way by which the presynaptic machinery acts is through RIMs, which recruit both voltage gated calcium channels and Munc13s, a family of proteins essential for making synaptic vesicles tethered at the active zone fusion competent (Südhof, 2012; Imig et al., 2014; Acuna et al., 2016). Bassoon regulates this core transmitter release machinery, at least at some synapses, by selectively recruiting the P/Q type of voltage gated calcium channels and by speeding up synaptic vesicle reloading to release sites during ongoing activity (Davydova et al., 2014; Hallermann et al., 2010; Mendoza-Schulz et al., 2014). In addition to regulating transmitter release, Bassoon and Piccolo maintain synaptic integrity by reducing the proteasome- and autophagy-mediated degradation of presynaptic molecules (Waites et al., 2013; Okerlund et al., 2017; Hoffmann-Conaway et al., 2020; Montenegro-Venegas et al., 2021). At the electron microscopy level, the multimolecular complex of presynaptic scaffold proteins manifests as a meshwork of filamentous structures termed the presynaptic particle web (Philips et al., 2001) or cytomatrix of active zones, i.e., CAZ (Cases-Langhoff et al., 1996; Garner et al., 2000; Dresbach et al., 2001).

Bassoon is a particularly large CAZ molecule, comprising 3938 amino acids in the rat, and 3926 amino acids in humans (tom Dieck et al., 1998). It shares 10 regions of sequence homology with Piccolo/Aczonin (Fenster et al., 2000; Wang et al., 1999). Light microscopy super-resolution studies and electron microscopy studies have revealed that Bassoon and Piccolo are oriented in a particular way at synapses, with their C-termini closer to the active zone plasma membrane than their N-termini (Dani et al., 2010; Limbach et al., 2011). Thus, Bassoon and Piccolo appear to be extended proteins with a parallel orientation at synapses, consistent with the assumption that they may represent some of the filamentous CAZ structures observed by electron microscopy.

Using recombinant Bassoon constructs (Dresbach et al., 2003) we previously imaged the incorporation of Bassoon into nascent synapses and its turnover at existing synapses (Shapira et al., 2003; Bresler et al., 2004; Tsuriel et al., 2006; Tsuriel et al., 2009). In the course of these studies, we also found that Bassoon– in addition to being a CAZ protein – is associated with the Golgi-apparatus, and that associating with the Golgi-apparatus is a prerequisite for the subsequent trafficking of Bassoon to synapses (Dresbach et al., 2006). Indeed, Bassoon, Piccolo and ELKS/CAST/ERC are all detected at the Golgi apparatus, and appear to exit the Golgi apparatus on transport vesicles that may carry CAZ material to synapses (Zhai et al., 2001; Maas et al. 2012). Unlike at synapses, the nanostructure of Bassoon at its second prominent subcellular localization, i.e., the Golgi-apparatus, has not been investigated. Here, we created a new generation of Bassoon constructs and determined their localization and arrangement at the Golgi-apparatus by stimulated emission depletion (STED) microscopy. We find that Bassoon is an extended molecule at the trans-Golgi-network (TGN) with its N-terminus closer to the TGN than its C-terminus.

## 3. Materials and Methods

### 3.1. Animals

Cells and tissues used in the study were obtained from bassoon gene trap (Bsn^GT^) (Hallermann et al., 2010); mouse strains backcrossed over more than 10 generations to C57BL/6N. Bsn^GT^ mice were obtained from Omnibank ES cell line OST486029 by Lexicon Pharmaceuticals, Inc. (The Woodlands, TX). All experiments were performed in accordance with the European Committees Council Directive (86/609/EEC) and approved by the local animal care committees (Landesverwaltungsamt Sachsen-Anhalt, Germany, and the State Government of Lower Saxony, Germany.

### 3.2. Antibodies

The following antibodies were used for immunocytochemistry (IC): mouse anti-Bassoon (1:500 ENZO life systems), rabbit anti-Piccolo(1:200 Synaptic systems), mouse anti-TGN38 (1 to 500BD-Transduction Laboratories), mouse anti-Syntaxin 6 (1:300 Abcam), chicken anti-GFP (1: 3000 Abcam), rabbit anti-GFP (1: 1000 Abcam), mouse anti-Synaptophysin (1: 1000 Sigma Aldrich), guinea pig anti-SHANK2 (1 to 1000 Synaptic systems), Nanobodies: RFP-Booster-Atto594 and GFP-Booster-Atto647 (1:300 Chromotek), Mouse AlexaFluor®647 (1:1000 for Epifluorescence microscopy/1:100 for STORM Invitrogen), Chicken Cy5.5 (1:150 for STORM Jackson/Invitrogen), Chicken and Rabbit Alexa Fluor®488 (1:1000 Invitrogen), ATTO-TEC dyes: Mouse AttoKK1212 and Mouse Atto594 (1: 100) and Abberior dyes: Mouse STAR 638, Mouse STAR 635p and Rabbit STAR 639 (1: 100).

### 3.3. Full-length Bassoon constructs

Full-length rat Bassoon constructs mRFP-Bsn-mEGFP, mEGFP-Bsn, Bsn-mEGFP, mRFP-Bsn, Bsn-mRFP, and mutant G2A-mRFP-Bsn-mEGFP were created in the ampicillin-resistant pCS2^+^ vector backbone and designed to include either an intramolecular mRFP/mEGFP tag and/or a mEGFP/mRFP fused to the C-terminus of Bassoon. The intramolecular tag was created by gene synthesis in a way that its coding sequence, after insertion into the HindIII site of rat Bassoon, was preceded by amino acids 1-97 of Bassoon and was followed by amino acids 95-3938 of Bassoon. We generated the G2A-mRFP-Bsn-mEGFP myristoyl mutant by inserting a point mutant at the second amino acid position, thereby replacing a glycine amino acid with alanine in the sequence of the full-length mRFP-Bsn-mEGFP construct. The plasmid called “CFP-Golgi” (containing amino acids 1–81 of ß-1,4glycosyltransferase) was originally purchased from Clontech.

Bassoon-mRFP was cloned by replacing mEGFP of the Bassoon-mEGFP construct with an mRFP tag using the Mlu and SpeI sites. Similarly, mEGFP-Bassoon was cloned by replacing mRFP-Bassoon by inserting a mEGFP, at the intramolecular 97^th^ amino acid position, by using the HindIII site. CFP-ß-1,4glycosyltransferase (amino acids 0—81), also known as CFP-Golgi was generously provided by also Craig Garner. Bassoon-mRFP was cloned by replacing mEGFP of the Bassoon-mEGFP construct with an mRFP tag from the Mlu-mRFP-SpeI-PUC vector (Clonetech). Both constructs were linearized via a Mlu and SpeI digestion, followed by a CIP dephosphorylation and ligated. The ligation mix was transformed into XL1 Blue competent cell and was grown on LB–ampicillin plates. Extracted DNA from the colonies grown on the plates were sequenced with Bsn_FW_1, Bsn_FW_3c, Bsn_FW_4, and Bsn_FW_5 sequencing primers to confirm colonies the successfully cloned and direction of insert in the construct. Similarly, we cloned mEGFP-Bassoon by replacing mRFP-Bassoon by inserting a mEGFP at the intramolecular 97^th^ amino acid position by using the mEGFP-HindIII-PUC vector (Clonetech) and a HindIII digestion followed by the same protocol as described for Bassoon-mRFP generation.

Bsn_FW_1 CTAATGGGAGGTCTATATAAG

Bsn_FW_2 AGCACTAGCTGGCGGCGGACA

mRFP-FW-3a GTAATGCAGAAGAAGACCATG

mRFP-Rev-3b CATGGTCTTCTTCTGCATTAC

Bsn_FW_3c GGGCTTCAAGTGGGAGCG

Bsn_FW_4 GGGCCAGGAGGAGACAGACG

Bsn_FW_5 GCTCCAAACCGGCAGCCAAAG

### 3.4. Primary hippocampal neuron cultures

#### Rat cultures

E19 rat hippocampi were dissected as previously described (Dresbach et al., 2006). Hippocampi were dissociated by a 20 min of trypsin treatment at 37°C and trituration. 50,000 dissociated neurons/cm^2^ were grown on poly-lysine coated coverslips (Sigma Aldrich) in Neurobasal medium enhanced with 2% B-27 and 0.5% L-Glutamine (Life technologies). Primary hippocampal neurons growing on either 12mm or 18mm coverslips, in 24-well or 12-well plates, respectively, were transfected with the calcium phosphate method at DIV3 and fixed, with 4% paraformaldehyde, for imaging of mature DIV15—30 transfected cultures. The protocol for which was performed as described previously^3^ (Figures 1—2, Suppl.1). Neurons were prepared for calcium phosphate transfection by replacing and saving the conditioned medium with 500µl (for 12mm coverslips) or 750µl (for 18mm coverslips) Optimem (Life Technologies) at 37°C and incubated for 20—30min. A DNA mix containing 7µg of DNA and 250mM CaCl_2_ was vortexed during the dropwise addition of 105µl of transfection buffer (274mM NaCl, 10mM KCl, 1.4mM Na2HPO4, 15mM glucose, 42mM HEPES, pH 7.06) and left to incubate for 20min at room temperature. 30µl per well of the DNA-Calcium phosphate mix was applied onto neurons and incubated further for 75min. We used three washes of pre-warmed Neurobasal medium to complete the transfection, reinstated the transfected neurons into their original conditioned medium, and incubated them at 37°C and 5% CO2 until the transfected cultures had matured.

**Figure 1.**
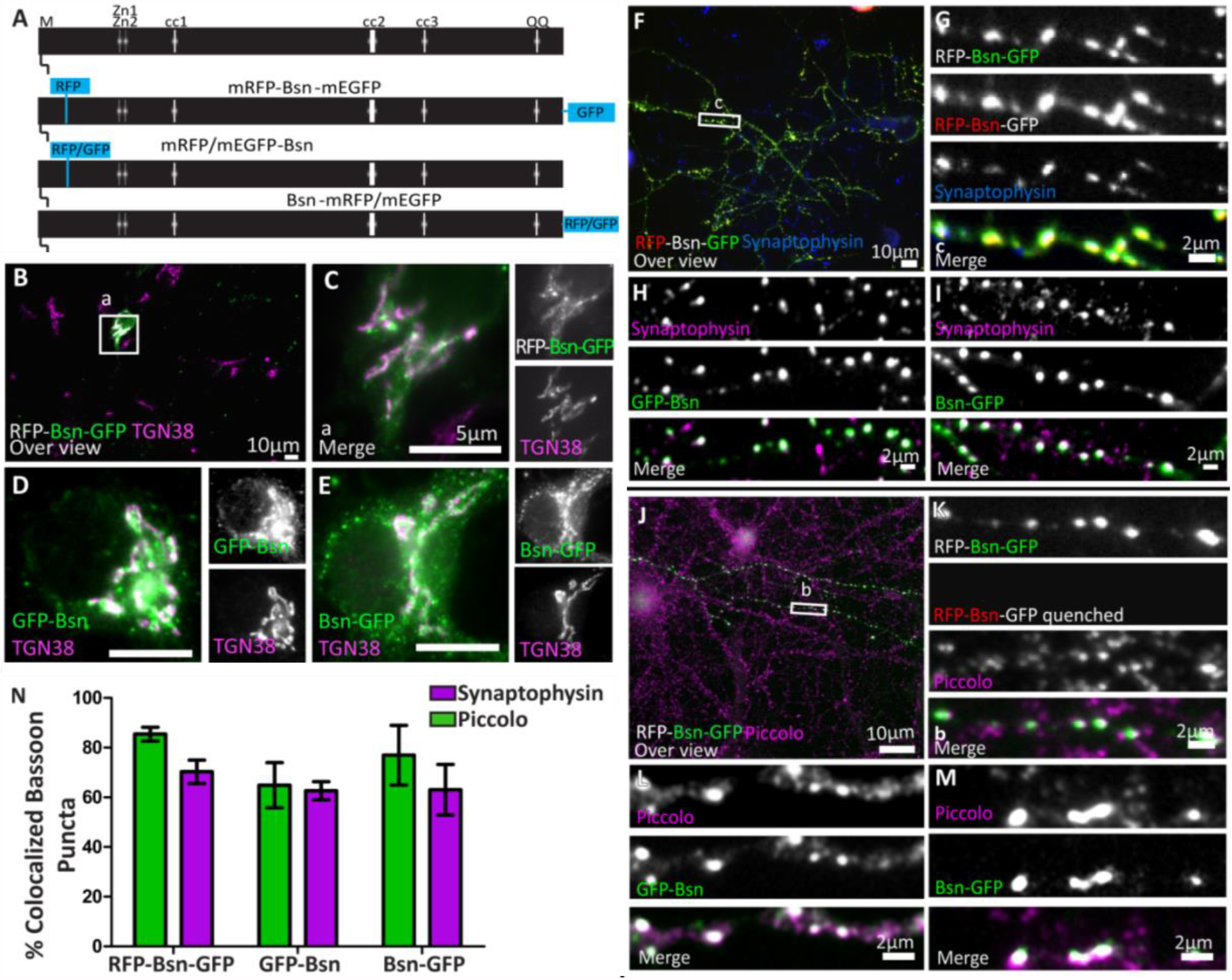
Full-length Bassoon constructs localize at the TGN in young neurons, traffick to synaptic sites, and are incorporated into the insoluble AZ scaffold of mature neurons. Panel **A** is a schematic diagram of full-length Bassoon sequence compared to the sequence of full-length double- and single-tagged (either mRFP / mEGFP tagged) Bassoon constructs where M stands for N-myristoylationsequence, Zn1 and Zn2 are the two zinc finger domains and cc1, cc2, cc3, are the three predicted coiled-coil regions. Immunostained DIV7 (**B—E**) and DIV14 (**F—M**) hippocampal neurons transfected with full-length double-and single-tagged Bassoon constructs with GFP, post a DIV3 lipofectamine transfection, are co-stained with the TGN38 (**B—E**), synaptophysin (**F—I**) and Piccolo (**J—M**) markers. Panels **B, F**, and **J** represent 40X over views of the transfections and **C, G** and **K** represent the zooms of their white square ROIs, respectively. Neurons in panels **B**,**C, J** and **K** were briefly fixed in cold methanol prior to normal fixation, to quench the RFP and GFP autofluorescence. Panels **J**—**K** are stained with TGN38 (purple) and a GFP antibody(green). Panels **J—M** are immuno-labeled for GFP antibody (green) and Piccolo (purple) after a 90 second treatment of 0.1%Triton X-100 and five minute methanol wash. **N** is the colocalization quantification of anti GFP immunofluorence of Bassoon for panels **F—M**; data are represented as mean ± SD, N=5 cells from two separate experiments for each quantification. Scale bars 10*μ*m (**B, F**, and **J**), 5*μ*m (**C—E**) and 2*μ*m(**G— I** and **K—M**).

**Figure 2.**
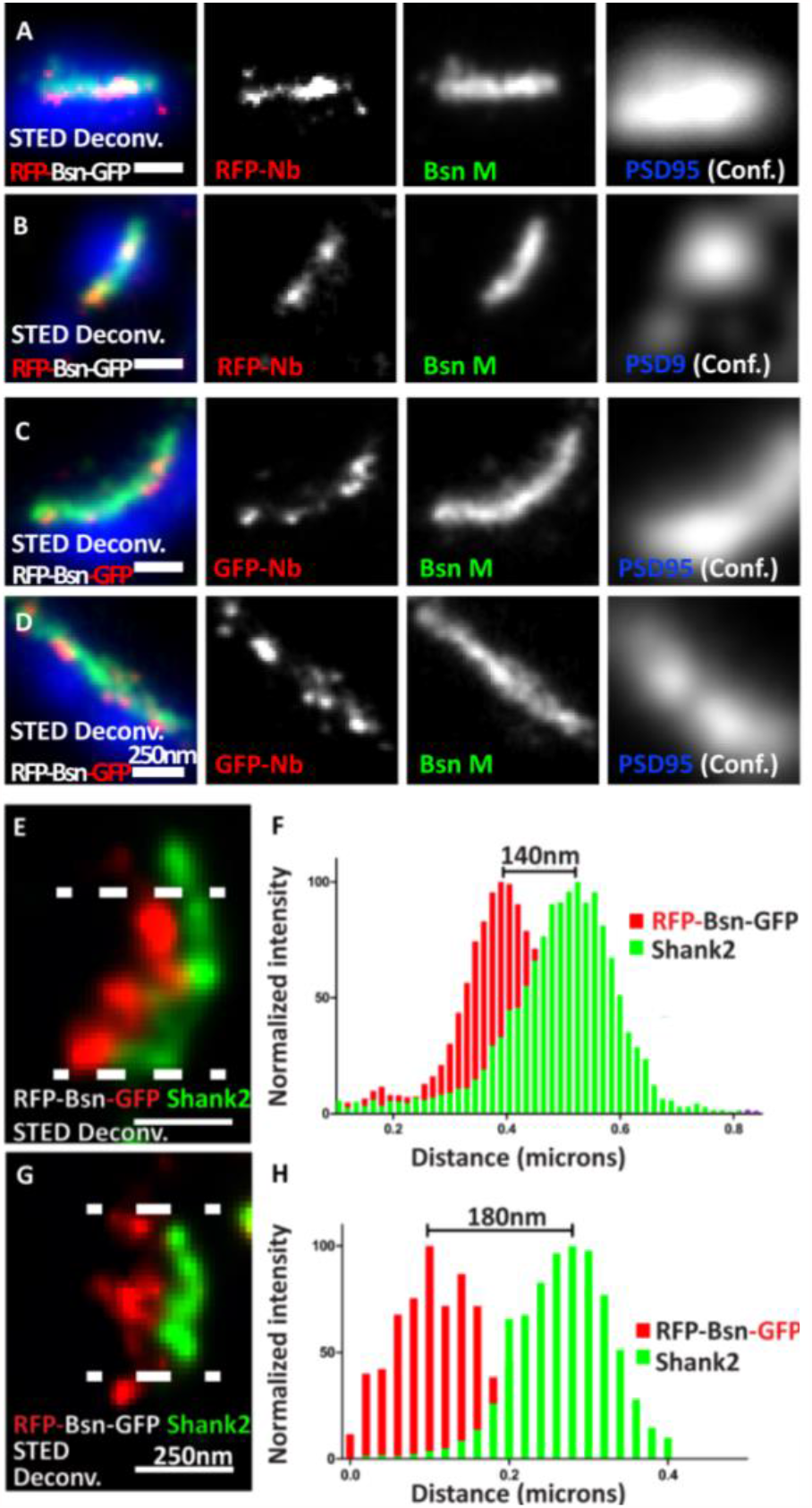
Visualizing the termini of full-length mRFP-Bassoon-mEGFP construct and its orientation with nanobodies, at mature synaptic sites. DIV21 mass hippocampal cultures transfected with the mRFP-Bassoon-mEGFP construct and imaged using two-color STED microscopy. The construct was visualized at synaptic sites with postsynaptic scaffold markers PSD95 and Shank2, using either the RFP-nanobody-Atto594 to visualize the RFP tag and the N-terminus of the Bassoon construct (**A, B**, and **G**) / the GFP-nanobody-Atto594 to visualize the GFP tag and the C-terminus of the Bassoon construct (**C—E**). Panels **A—D** are two-color-STED deconvolved (Deconv.) images of triple color stainings that label the endogenous presynaptic Bassoon signals with a traditional monoclonal antibody, nanoclusters of nanobody signals within the endogenous presynaptic Bassoon signals, and postsynaptic scaffold marker PSD95 (in confocal mode). Panels **E** and **G** show the localization of nanobody labelled C- and N-termini of mRFP-Bassoon-mEGFP and Shank2 in side-view images of its synapses. Distribution of localization points within a 350nm thick line profiles at the center of the synapse (as shown by the area within the dashed lines) were measured, fit with gaussian distributions and are plotted in panels **F** (C-terminus of tagged Bassoon and Shank2) and **H** (N-terminus of tagged Bassoon and Shank2). The distance between centroids of the two Gaussians defines the Bassoon-Shank2 distances. Scale bars 250nm (**A—G**).

Mouse cultures were prepared as described (Montenegro-Venegas et al., 2021). Briefly, P0-P1 wild type mice cortexes were dissected to generate the feeder layer of a sandwich culture. Hemispheres of cortexes with their meninges removed were chopped up in 4.5 ml of HBSS and incubated for 15 minutes at 37°C in 2.5% trypsin (without EDTA). These pieces were then washed in HBSS and dissociated in glia medium that consisted of 90% plating medium and 10% DNAse (Invitrogen). Both hemispheres of one brain were dissociated in 1 ml glia medium and plated in 10 ml of plating medium that was identical to rat primary culture plating medium. The medium was changed every 4—5 days, and the confluent glia were trypsinated, washed in HBSS, and 5 ml of the glia were plated on a 6cm dish. Two P1 Bsn-/- knockout mice and two wildtype Bsn+/+ littermates were prepped into dissociated primary hippocampal neurons following the same protocol as was used for rat primary culture. 100 ml of 5000 hippocampal cells were plated on coated 18 mm round glass coverslips. These coverslips were first incubated for 1 hour at 37°C and 5% CO2 and then transferred, neurons facing down, onto dishes containing the feeder layer of glia and 5 ml of culturing medium (94% Neurobasal, 2% Glutamax (Invitrogen), 2% B27, 1% NaPyr (0.1M), 1% Pen/Strep (0.1M). These coverslips were left to grow at 37°C and 5% CO2 and were treated with 2 µl Ara-C (Sigma) on day in vitro 1 (DIV1) and DIV3 to prevent glia overgrowth and were fed once a week with an exchange of 1 ml of fresh culturing medium to maintain optimal growth of the culture.

### 3.5. Transfection

To visualize Golgi association of Bassoon constructs in hippocampal neurons (Figures 1, 3—6), we applied the Lipofectamine transfection method on DIV5/6 neurons and fixed them on DIV6/7 neurons with 4% paraformaldehyde, as previously described. Briefly, a conditioned medium of neurons, on 12mm coverslips, was exchanged with 500ml pre-warmed Neurobasal medium, containing 2% B-27 and 1% of 2mM L-Glutamine, saved and incubated along with the neurons for 20—30min. A Lipofectamine solution and a DNA solution of 25µl Optimem/well (Life Technologies) with 1µl of Lipofectamine 2000/well (Invitrogen) and 1µg of plasmid DNA/well were prepared and mixed after 10min room temperature incubation. The Lipofectamine-DNA mix was further incubated for 20min at room temperature; 50µl/well of the solution was dropwise applied on the neurons and incubated for 75min in 37°C and 5% CO2 conditions. The transfection was completed after three pre-warmed Neurobasal washes and reinstating the transfected neurons in their conditioned medium at 37°C. These neurons were fixed the next day for 20min in cold 4% paraformaldehyde solution before immunocytochemistry was performed.

**Figure 3.**
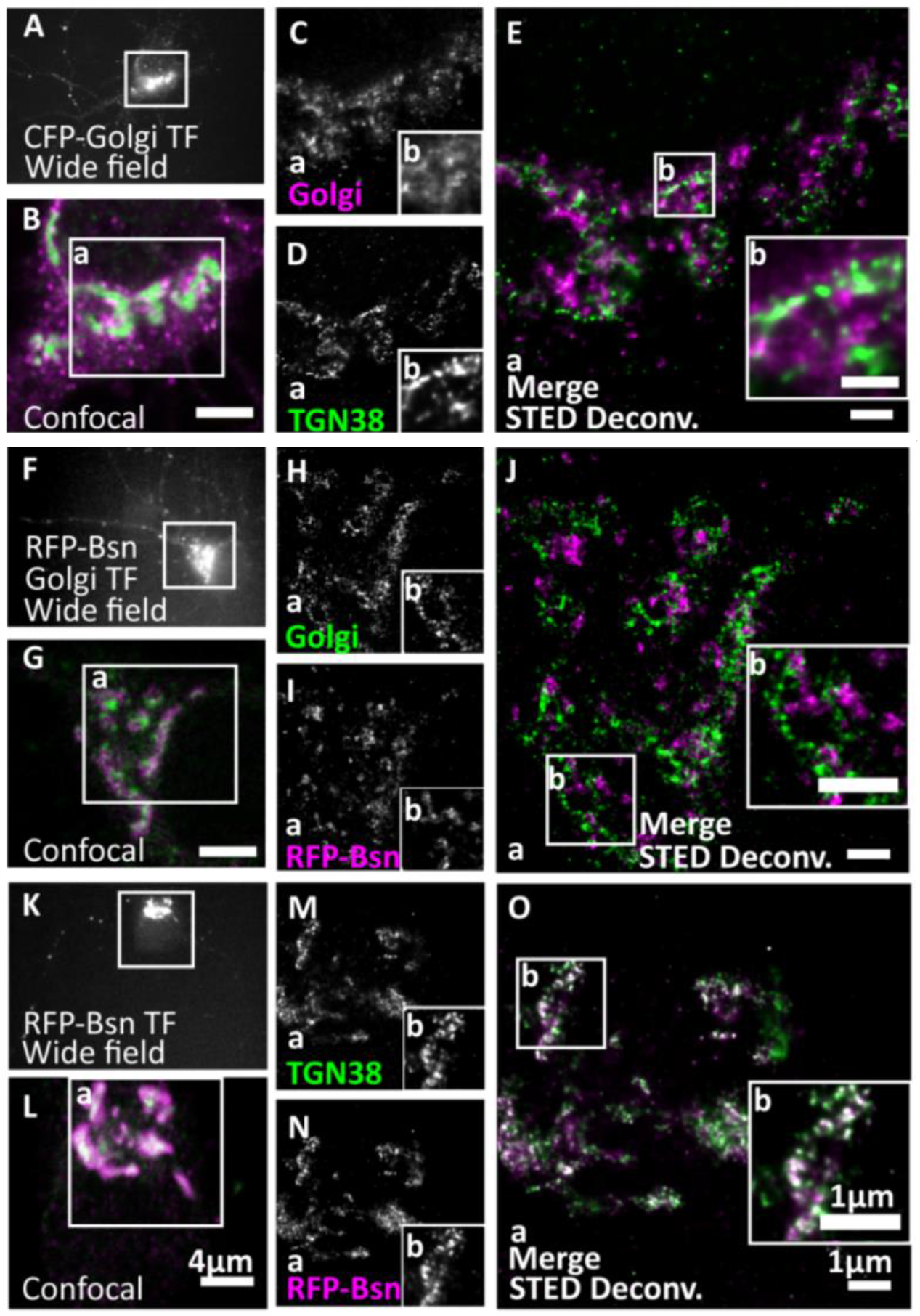
Full-length Bassoon localizes to the trans-Golgi network, not the trans-Golgi sub-compartment. DIV7 hippocampal neurons were transfected with CFP-Golgi (*trans*-Golgi sub-compartment maker), full-length single-tagged mRFP-Bsn construct, and immunostained using GFP and/or RFP nanobodies against tagged constructs and from **A—E** with TGN38 (trans-Golgi network maker). Two-color STED images of both Golgi sub-compartment markers (**A—E**), CFP-Golgi and RFP-Bsn constructs (**F— J**), and RFP-Bsn at TGN38 (a *trans*-Golgi Network maker) (**K—O**). **A, F, K** show wide field overview of transfected construct, **B, G, L** the confocal zooms of the soma, insets **a** reflect the single channels and merged full STED deconvolved (Deconv.) images of **C—E, H—J**, and **M—O**, while insets **b** represents the zooms of STED images. Scale bars 4μm (**B, G** and **L**) and 1μm (**E, J** and **O)**.

**Figure 4.**
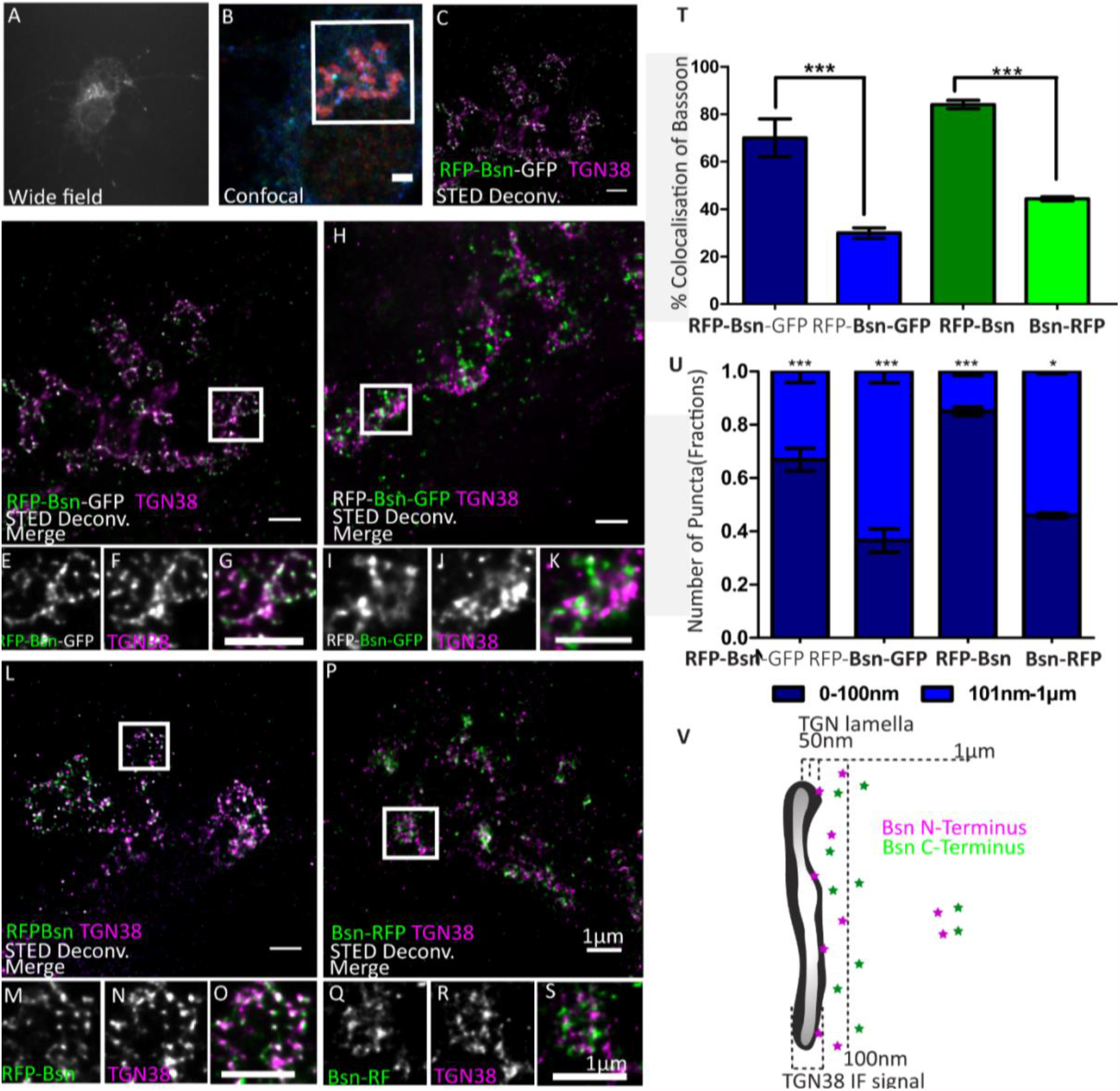
Orientation double-and single-tagged Bassoon constructs at the *trans*-Golgi network marker TGN38. DIV7 hippocampal neurons transfected with double-(**A—K**) and single-tagged (**L—S**) full-length Bassoon are immunostained with the *trans*-Golgi network marker TGN38 and an RFP-nanobody-Atto594 to visualize the RFP tag (**A—G, L—O** and **P—S**) / a GFP-nanobody-Atto594 to visualize the GFP tag (**H—K**). The experimental schematic (panels **A—C**) demonstrates the acquisition of two-color STED images wherein the wide field overview of the transfected neuron (**A**), its corresponding confocal over view of the soma, with GFP autofluorescence in blue, (**B**) and a 10μmX10μm inset scanned in STED mode to visualize the constructs’ N—terminus (**C, D** and **L**) and C—terminus (**H** and **P**). Merged and single channel views of the zoom images of **E— G, I—K, M—O**, and **Q—S** are representations of the white ROIs in the STED images **D, H, L** and **P** respectively. **T** represents the colocalization quantification, and **U** represents the allocation quantification at and away from TGN lamella i.e. 0—100nm or 101nm—1μm, respectively. Data are represented as mean ± SD, N=10 cells from two separate experiments, statistically tested with a one-way ANOVA with the Tukey’s multiple comparison’s post-hoc test \*\*\**p* ≤ 0.001. **V** is a Schematic representation of the proposed distribution of N- and C-terminal Bassoon immunosignals at the TGN. Scale bars 2μm (**B**) and 1μm (**C—S**).

**Figure 5.**
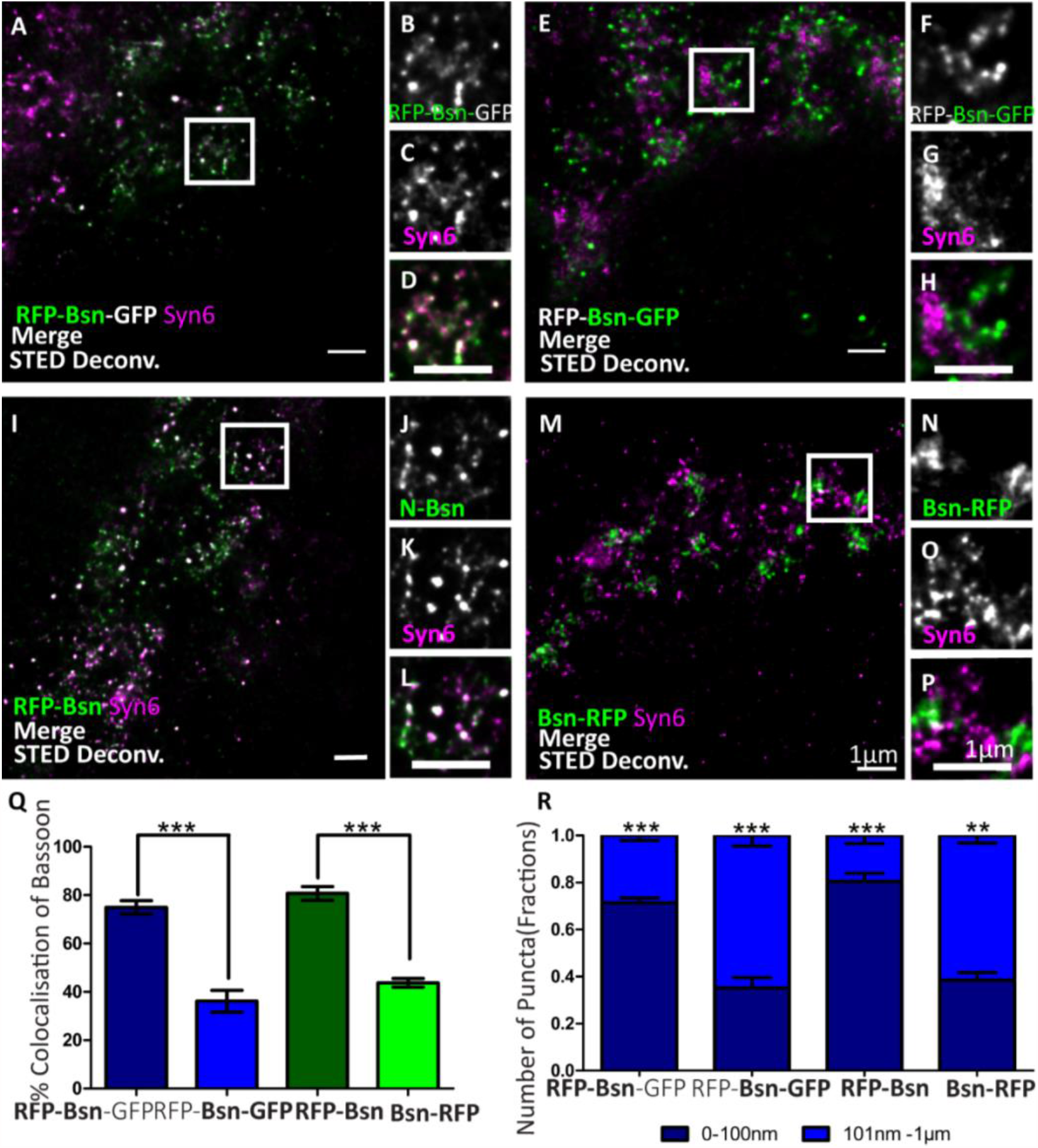
Orientation double-and single-tagged Bassoon constructs at the *trans*-Golgi network marker Syntaxin 6 (Syn6). Transfected DIV7 hippocampal neurons immunostained for either one or both termini of full-length Bassoon with RFP-nanobody-Atto594 / GFP-nanobody-Atto647 and the Syn6 marker. Two-color, deconvolved, 10*μ*mX10*μ*m STED images and their ROI zooms, show double-tagged and single-tagged Bassoon constructs in panels **A—D** and **I—L** (of which the N–termini of constructs were imaged) and panels **E—H** and **M—P** (of which the C–termini of the constructs were imaged), respectively. Graph **Q** quantifies the amount of Bassoon colocalization with Syn6 and graph **R** quantifies the Bassoon signal allocations at and away from TGN38 lamella i.e., 0—100nm or 101nm—1*μ*m, respectively. Data are represented as mean ± SD, N=10 cells from two separate experiment, statistically tested with a one-way ANOVA with the Tukey’s multiple comparison’s post-hoc test **p ≤ 0.01 and ***p ≤ 0.001. Scale bars 1*μ*m (**A—P**).

**Figure 6.**
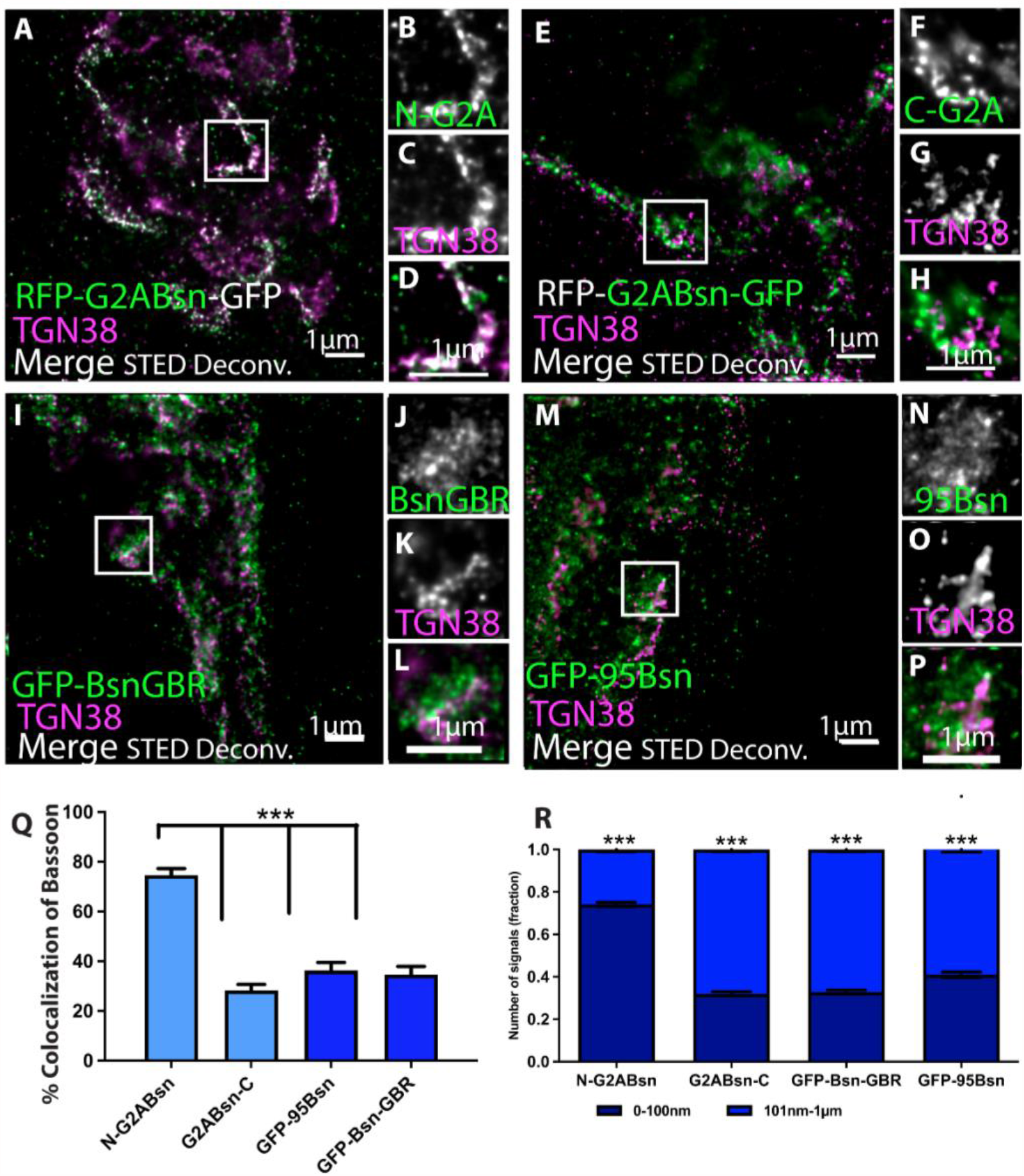
Orientation of deletion constructs of Bassoon at the *trans*-Golgi network. DIV7 hippocampal neurons transfected with tagged Bassoon deletion constructs a) myristoyl group deficient G2A-mRFP-Bsn-mEGFP (**A—H**), Bassoon’s N–terminus, i.e., (95-3938) 95-Bsn construct (**I—L**), and Bassoon’s N– and C–termini (2088-2038), i.e., Bsn-GBR construct (**M—P**) were visualized with a GFP-nanobody-Atto647 / RFP-nanobody-Atto594 and TGN38 marker. Insets of the two-color STED deconvolved images in **A, E, I** and **M** are represented in panels **B—D, F—H, J—L**, and **N—P**, respectively. Graph **Q** and **R** quantifies the colocalization and signal allocations of Bassoon at the TGN38, respectively. Data are represented as mean± SD, N=10 cells from two separate experiment, statistically tested with a one-way ANOVA with the Tukey’s multiple comparison’s post-hoc test \**p* < 0.05 & \*\*\**p* ≤ 0.001. Scale bars 1μm (**A—P**).

### 3.6. Immunocytochemistry

Primary cultures of hippocampal neurons were washed multiple times after the paraformaldehyde fixation, blocked for 20 min with the primary antibody-buffer (10% FBS, 5% sucrose, 2% albumin, 0.3% Triton X-100 in 1× PBS) at room temperature, and stained with the primary antibodies, diluted in the primary antibody-buffer, overnight at 4°C. Following multiple washes, secondary antibody dilutions were prepared in the secondary antibody-buffer (0.3% Triton X-100, 5% sucrose, and 2% albumin in 1× PBS) and applied on the coverslips for 1 hour, in darkness, at room temperature. Three washed of 1xPBS and one of distilled water were performed on the coverslips before being mounted on slides with DABCO-mowiol (Calbiochem) and left to dry overnight.

#### 3.7.1. Microscopy and analysis Epifluorescence microscopy

An inverted Zeiss fluorescence microscope (Observer.Z1) with a Photometrics CoolSnap HQ2 camera was used to image samples at a magnification of 40X and 63X. The following filters from AHF were used: F46-000 for DAPI, F46-002 for GFP and Alexa 488, F46-004 for Atto594 and Alexa 546 dyes, and F46-006 for AttoKK1212 and Alexa 647. Exposure times of 500ns for F46-002 and F46-004 filters and 1000ns for the F46-006 filter were applied. The images were processed with the Image J software (NIH) (imagej.nih.gov/ij/) to generate scale-bar inserted RGB merged TIFF files for further analysis with Imaris MeasurementPro software. Images in Figure 1 and Supplementary 1 were adjusted for brightness and contrast in Image J, where needed, calculated, and stamped with a suitable sized scale bar.

#### 3.7.2. Colocalization analysis for conventional epifluorescence images

Merged multi-channel 40X light microscopy images were analyzed using MetaMorph Offline Version 7.7.0.0 (Molecular Devices, Inc.). A threshold is set for each channel followed by the generation of a mask for all channels, in three areas of size 25 pixels long (representing 2 µm in the sample) and 4 pixels wide, per image. These area masks were then overlaid in the arithmetic tool and divided to generate a third mask containing only the population of fluorescence signals in the mask that do colocalize. The amount of bassoon colocalized is represented as a percentage of the bassoon colocalized population divided by the total bassoon population.

#### 3.7.3 STED microscopy

STED images in Figures 2—6 were acquired on a custom-built two-color STED microscope that includes a 1.4 NA 100× objective (PL APO HCX 100x 1.4-0.7 Oil, Leica Microsystems, Wetzlar), and a 775nm STED laser (ELP-5-775-DG, IPG Photonics Corporation, Oxford, MA, USA)^4^. The dyes were excited at wavelengths of 470nm, 595nm, and 640nm, while the fluorescence was detected with avalanche photodiodes from 500-550nm, 600-640nm, and 660-720nm, respectively. Corroborative confocal images were acquired using a LED illumination source, a monochrome filter, and a camera (DMK41 AU02, The Imaging Source). The LED illumination source for such overview images was manually installed every session, wherein a 590nm LED was installed with the 700/60 fluorescence filter, and 640RDC dichroic filter or a 490nm LED was installed (upon requirement) with the 450/60 fluorescence filter in the camera path. Images were taken at 300—700 mW STED power, 4µW excitation power, dwell time of 30—100ms and a pixel size of 10nm. The resolution regularly obtained for Atto594 antibodies was 25-40nm and for AttoKK1212 antibodies was 20-35nm (at 300mW and 700 mW STED-power, respectively). Figures 2-6 were acquired on the Abberior QuadSCAN two-color STED microscope. The setup was equipped with a pulsed 775nm STED lazer and two pulsed excitation lazer-sources at 594nm and 640nm integrated into an Olympus IX83 microscope. The setup also included a 100× 1.4NA objective; a 4-color LED illumination source, a gated avalanche photodiode (APD), and a wide monochrome field. A pixel size of 10 nm, dwell time of 3ms, and three-line accumulations were applied to the images. All STED images were acquired with the ImSpector Software^5^ (Max Planck Innovation) and were processed using the Richardson—Lucy deconvolution function. In combination with the deconvolution processing, a 2D Lorentz function, that fits the full width at half maximum (FWHM) fitted of the point spread function of each individual image to the resolution estimate was used. These images were then analyzed using Imaris MeasurementPro. Images for figures were adjusted for brightness and contrast with Image J software (NIH).

#### 3.7.4 Quantitative analysis of the probability and amount of colocalization and the distribution of Bassoon constructs at the Golgi

Merged TIFF epifluorescence images or STED images were analyzed using Imaris MeasurementPro 8.1(Bitplane AG.) software to ascertain the probability of colocalization (Pearson’s correlation coefficient), amount of colocalization, and distribution pattern of signals within the images.

Soma Images in Figures 3—6 were analyzed within a ROI, generated by a free-hand drawn mask, to exclude any signals in the image that was in the nucleus, outside the cell soma, or in a neighboring neuronal process. Images were analyzed for the probability of colocalization through the ImarisColoc module. The integrated Costes P-Value approximation plugin^7^, within the ImarisColoc module, first generates automated thresholds for all the channels of all the images, which are subsequently used to calculate the colocalization Pearson’s correlation coefficient constants in the picture. The amount of colocalization and distribution pattern of signals was calculated after Imaris Spots generation. Imaris Spots, a built-in spot detection algorithm, was used to generate objects for every punctuate signal, for all channels, in the images. These spot objects were defined by the automated intensity threshold value for each channel (calculated by the Costes P-Value approximation), signal diameter size range (10nm— 160nm for STED images and >200nm for light microscopy images), and an automated splitting of signal clusters (defined as >120nm for STED images and >400nm for light microscopy images), for each channel.

The amount of colocalization was calculated using Spots Colocalize, a MATLAB extension in the Imaris Spots module, at a distance threshold of 0—100nm (for STED images) and 0—350nm (for light microscopy images) from the spot centers.

The distribution pattern of AZ protein signals at the Golgi was extracted by applying the distance transformation MATLAB extension^8^, from the Imaris XT module to the spots object information generated via the Imaris Spots module. This distance transformation extension was applied on the Golgi label channel transforming the Golgi voxel intensity data into spot coordinates. These spot coordinates were then used to calculate the shortest distance of each AZ protein spot object to the object border of a Golgi-label spot. The data for the shortest distances between all the AZ protein signals and the border coordinates of the Golgi-label, and the total number of AZP signals within 0—100nm or 101—1000nm distance ranges was extracted from the statistics of the Imaris Spot module, statistically tested, and graphically represented with GraphPad Prism.

The distribution pattern of Bassoon constructs in Figure 7A was generated in Python and graphically visualized as violin plots, with a signal size upper limit of 600nm; to clearly represent the distribution and permit visualization of the median and interquartile ranges of the data. The average distance of each cell per set, in Figure 7B, was extracted from the Imaris Spots statistics and limited to the 220nm cut-off were plotted in GraphPad Prism.

**Figure 7.**
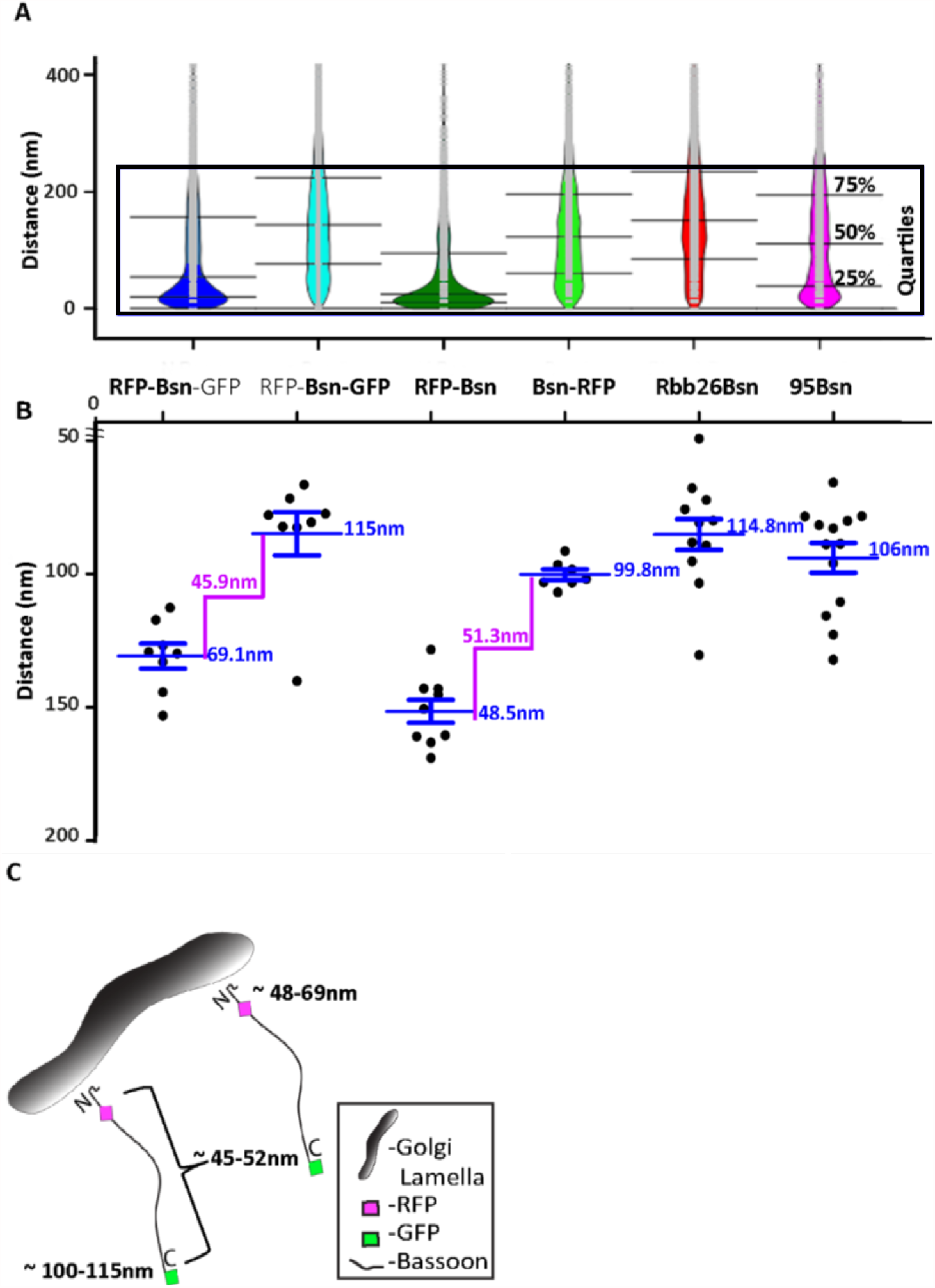
Distribution and average distances of all full-length and mutant Bassoon signals from the TGN38 reference maker. Bean plots in panel **A** display all signal distributions of single- and double-tagged full-length and deletion constructs of Bassoon from their nearest TGN38 marker. The light grey lines within each individual bean plot show individual observations of Bassoon signals. The blue ROI represents the 220nm range that corresponds to 75% of the total Bassoon signals oriented around the TGN lamella (**A**). The average distances of the Bassoon tags of all five constructs, within the 200nm distance range, are plotted in **B. C** is a schematic diagram that shows the average distance N and C-termini of full-length Bassoon molecules possess at the TGN lamella.

Statistical analysis and representation of all resultant data were prepared in GraphPad Prism 5.02. Data are presented as mean ± SEM, and statistical differences were considered significant, strongly significant, and extremely significant at respective p values of *p < 0.05, **p ≤ 0.01, and ***p ≤ 0.001. In a two-step statistical testing protocol, first, a one-way ANOVA test, with a post-hoc Tukey’s multiple comparisons test, was performed, and for every significant difference observed between two relevant groups, an additional two-tailed, unpaired Student’s t-test, with different variances, was also performed to reveal the same significant difference.

#### 3.7.5 Quantitative analysis of distances between Bassoon’s termini and the post synapse

All acquired, or rendered images were processed and visualized using ImageJ (imagej.nih.gov/ij/) and ImSpector software (Max-Planck Innovation) (Figure 2) or Daxviewer software (Mark Bates) (Figure 3). Line profiles were measured with ImageJ software along a 350nm thick line profiles. Inter-peak Bassoon-SHANK2 distances were determined after fitting a Gaussian distribution in GraphPad Prism.

To factor out the effect of the varying number of signals counted per size of hand-drawn mask in each image, the area in µm3 of the mask used, was divided by the signals measured per image.

## 4. Results

### 4.1. Characterizing second generation full-length Bassoon constructs

We first aimed at improving three features of recombinant Bassoon:

a. Faithful expression of the full-length protein including its C-terminal tag: when expressed in neurons, Bassoon-1-3938-EGFP, in addition to producing punctate synaptic fluorescence signals, also produces diffusely distributed green fluorescence, presumably resulting from soluble EGFP or a soluble C-terminal fragment of Bassoon with the EGFP-tag attached (Dresbach et al., 2003). We realized serendipitously that this diffusely distributed green fluorescence also occurred when the EGFP coding sequence was attached out of frame to the 3’ end of Bassoon, suggesting that a cryptic ribosomal entry site exists somewhere near the 3’-region of Bassoon or in the linker located between Bassoon and EGFP. To prevent translation of the C-terminal EGFP-tag we changed the linker sequence and removed the start ATG from the EGFP coding sequence.
b. The accessibility of its N-terminus: Bassoon contains a functional consensus site for N-myristoylation (Dresbach et al., 2003), so N-terminal tags might impair N-myristoylation. Ideally, a tag designed to locate the N-terminal region of Bassoon should be placed downstream of the N-myristoylation consensus site. To leave this consensus site unaffected, we placed either RFP, CFP or GFP as an intramolecular tag 97 amino acids downstream of the N-terminus of Bassoon, using an endogenous HindIII site in the rat Bassoon cDNA. We will refer to these tags as “intramolecular N-terminal” tags, to highlight both of their features, i.e., leaving the very N-terminus intact, and placed close to it.
c. Its tendency to aggregate: Bassoon may form homodimers, and heterodimers with Piccolo (Maas et al., 2012). Tags with an inherent capacity to dimerize could cause aberrant oligomerization and generate non-functional aggregates of Bassoon. To prevent this, we used monomeric fluorescent proteins, including RFP, CFP and the A207K variant of EGFP. We use the term “EGFP” when referring to previously generated constructs, which harbor standard EGFP, and we use the term “GFP” when referring to new constructs, which harbor the monomeric variant. A schematic synopsis of these new constructs is presented in Figure 1A.

To test the new constructs, we transfected dissociated rat hippocampal cultures with the new full-length single-tagged and double-tagged Bassoon constructs on day 3 after plating (day in vitro 3; DIV 3). We characterized the subcellular localization of these Bassoon constructs in young (DIV5) and mature (>DIV13) neurons (Fig 1.), by immunostaining fixed cultures using a single-domain antibody (nanobody) directed against RFP, a polyclonal antisera directed against GFP, and monoclonal or polyclonal antibodies directed against markers for subcellular structures. The extended characterization of all single-tagged second-generation Bassoon constructs can be found in supplementary Figure 1.

In the soma of young neurons, all constructs were readily detected at the Golgi apparatus, labeled by the trans-Golgi-network (TGN) transmembrane protein TGN38 (Figure 1B—E). Previous reports observed similar localizations of endogenous and recombinant Bassoon signals in young neurons (Dresbach et al., 2006; Maas et al., 2012). To test the targeting of these constructs to synapses and their incorporation into the CAZ matrix, we analyzed their localization in mature neuronal cultures. In DIV14 neurons immunostained for the tags and the synaptic vesicle marker synaptophysin or the CAZ marker Piccolo, none of the constructs showed the diffusely distributed green fluorescence associated with the first-generation Bassoon-EGFP (Dresbach et al., 2003), and all of the constructs accumulated at synaptic sites (Fig. 1F-N). The degree of colocalization with synaptophysin was 70.25% (± 18.23% SD) for the dually tagged construct, 66.7% (± 12.9% SD) for mRFP Bassoon, and 62.64% (± 6.37% SD) for Bsn-mRFP (Figure 1F—I,N and supplementary Figure 1). Likewise, these Bassoon accumulations colocalized with the core CAZ scaffold protein Piccolo (Figure 1J—N and supplementary Figure 1), further corroborating their localization to synapses. This was true for the dually tagged (85.4% ± 18.42% SD) and the single-tagged Bassoon constructs (69% ± 5.6% SD for mRFP-Bsn and 76.97% ± 10.16% SD Bsn-mRFP).

Active zone proteins become resistant to Triton X-100 extraction once they became incorporated into the CAZ scaffold (Dresbach et al., 2003). When we applied a 0.1%Triton X-100 extraction to live neurons, followed by fixation and immunostaining for the tags and for endogenous Piccolo, we found that the synaptic accumulations of the recombinant proteins were indeed preserved. Colocalization with Piccolo was 85.43% (± 18.42% SD) for the dually tagged Bassoon, 64.85% (± 15.74% SD) for GFP-Bsn, and 76.97% (± 24.01% SD) for Bsn-GFP). The resistance of the recombinant Bassoon to the Triton X-100 treatment is indicative of their successful incorporation into mature active zone scaffolds (Figure 1 J—N).

### 4.2. Visualizing the orientation of full-length Bassoon constructs with nanobodies and super-resolution imaging

We then employed these constructs to study recombinant Bassoon by STED nanoscopy. To take full advantage of super-resolution microscopy, we used camelid antibodies, called nanobodies, to detect the tags. These anti-mRFP and anti-mEGFP nanobodies are small (1.5 nm x 2.5 nm) single-domain molecules derived from one heavy chain of an alpaca IgG antibody (Hamers-Casterman et al., 1993). They are designed to identify a single epitope on the tertiary structure of mRFP and mEGFP fluorophores (5nm diameters). These nanobodies were pre-coupled to two molecules of organic ATTO-TEC dyes, each 2—3nm in size. The mRFP nanobody was coupled to ATTO594, the mEGFP nanobody was coupled to ATTO647. Compared to traditional primary and secondary antibodies, which create a 30 nm labeling distance from the epitope site, the nanobody-ATTO dye complex generates a three times smaller label cloud around the tags (Wildanger et al., 2009).

Our microscopy setup allowed us to record triple-immunofluorescence images, with two dyes recorded in STED mode at a resolution of 20 nm, and a third dye recorded in confocal mode. In the first set of experiments, we analyzed cultures transfected with dually tagged Bassoon, asking three questions: 1) Do the nanobodies allow for detection of the construct in STED mode? 2) Can we spatially resolve the N- and the C-terminus of the dually tagged construct? 3) If so, is the recombinant construct oriented as predicted, i.e., with the C-terminus closer to the active zone than the N-terminus (Dani et al., 2010)?

Both the RFP-nanobody and the GFP-nanobody produced line-shaped or crescent-shaped signals (Fig. 2A-D), as expected for the appearance of active zone associated proteins at super-resolution (Dani et al., 2010). To make sure that what we analyzed represented synaptic Bassoon, we only analyzed signals fulfilling two criteria: they had to be line-shaped or crescent-shaped, suggesting that they represent side-view synapses, and, in addition, they had to colocalize with postsynaptic marker PSD95 immunofluorescence recorded in the confocal mode, corroborating that these signals represent synaptic Bassoon (Fig. 2A-D). Comparing the nanobody-signals to signals produced by a conventional monoclonal antibody directed against amino acids 756-1001 of Bassoon and detected by secondary antibodies revealed that the nanobody signals and the indirect immunofluorescence signals colocalized when analyzed by dual color STED microscopy. In addition, the nanobody signals produced sub-clusters of fluorescence within the lines and crescents (Fig. 2A-D). This is consistent with the assumption that their small size and their direct coupling to the fluorescent dyes allows for greater detection precision compared to indirect immunofluorescence with a conventional antibody. Overall, these data indicate that the RFP- and GFP-nanobodies can localize the N- and C-terminus of recombinant Bassoon molecules within the CAZ of mature neurons, and that the spatial precision of detection at least equals, and may exceed, the spatial precision provided by indirect immunofluorescence using a conventional antibody.

The orientation of endogenous Bassoon at synapses in brain sections was previously analyzed using conventional antibodies imaged by Stochastic Optical Reconstruction Microscopy (STORM) (Dani et al., 2010, Held et al., 2020). These studies used the monoclonal antibody to detect amino acids 756-1001 and/or a polyclonal antibody against the C-terminal 330 amino acids of Bassoon. They showed that Bassoon molecules possess an extended conformation within the CAZ scaffold and are oriented with their C-termini closer to the synaptic cleft than their N-termini. Using our dual color STED setup, we tested whether our constructs adopt a similar orientation in cultured neurons. We found that they indeed possess a similarly extended conformation, with the C-terminus 140 nm and the N-terminus 180 nm from the postsynaptic scaffold marker Shank2 (Fig. 2E-H). These constructs, in combination with nanobodies and STED microscopy, therefore, form an effective toolbox to visualize the nanoscopic localization and orientation of recombinant Bassoon in neurons.

### 4.3. Visualizing the orientation of new full-length Bassoon constructs at the Golgi-apparatus

How Bassoon is arranged at subcellular sites other than synapses has not been investigated by nanoscopy thus far. Bassoon may form primordial scaffolds at the level of the Golgi-apparatus (Dresbach et al., 2006; Maas et al., 2012). To arrive at a more comprehensive understanding of the possible arrangement of Bassoon, we used STED microscopy to determine the localization and orientation of our new constructs at the Golgi-apparatus of immature neurons, at DIV7.

We began by testing which Golgi compartment recombinant Bassoon is located to. The Golgi compartment is polarized across the flat stacks of cisternae/lamellae that form sub-compartments at its *cis* side, which receives materials from the endoplasmic reticulum (ER), and the Trans Golgi (TG) side, which sends them forward to their destinations. Additional vesicular-tubular structures both on the *cis* and *trans* ends of the Golgi stack, are known to form specialized compartments for cargo sorting at the entry and exit sides of the Golgi. The *trans*-Golgi network (TGN) is a post-Golgi compartment, following the TG sub-compartment that is involved in complex cargo sorting mechanisms (Griffiths andSimons, 1986; Mellman and Simons, 1992). We identified the TG sub-compartment and its neighboring compartment the TGN separately and compared the extent of co-localization of Bassoon at these two substructures in the soma of cultured neurons. To label the TG sub-compartment, we expressed a CFP-tagged construct (named CFP-Golgi) containing the cytoplasmic domain of ß-1,4-galactosyltransferase-11, which labels the last Golgi cisterna of the trans-Golgi compartment (Wittenmayer, 2014). To label the TGN, we immunostained for the marker TGN38. We detected recombinant constructs using the nanobodies, and we detected TGN38 using conventional primary and secondary antibodies. STED imaging revealed that the signals for the TG sub-compartment and TGN can be resolved and occupy different localizations in the soma (Fig. 3 A-E). mRFP-Bsn displayed virtually no colocalization with CFP-Golgi labeled TG lamellae (Fig. 3 F-J), but colocalized extensively with TGN38 (Fig.3 K-O), indicating that this construct is more closely associated with the TGN lamella than with the TG sub-compartment lamellae.

We then sought to determine the localization – and possibly the orientation – of Bassoon at the TGN in more detail. We first imaged the double-tagged Bassoon construct (RFP-Bsn-GFP) in somas of DIV7 neurons labeled with TGN38. The intramolecular N-terminal tag of the construct colocalized more extensively with TGN38 than its C-terminal tag (Fig. 4A-K). On average, 67.5% (±12.5% SD) of the signals coming from the intramolecular N-terminal tag, but only 29.8% (±7.7% SD) of the signals coming from the C-terminal tag colocalized with TGN38 (Figure 4T). To rule out that the difference could be caused by different avidities of the RFP- and the GFP-nanobodies we transfected neurons either with RFP-Bsn or with Bsn-RFP and detected the tags of both constructs using the RFP nanobody. Again, the intramolecular N-terminal tag colocalized more extensively with TGN38 than the C-terminal tag (Fig. 4L-S). In particular, the intramolecular N-terminal RFP showed 84.8% (±4.6% SD) colocalization while the C-terminally located RFP showed 44.4% (±2.4% SD) colocalization (Fig. 4T). Thus, both the dually tagged and the single tagged constructs reveal a closer apposition of the N-terminal region of Bassoon with TGN38 than its C-terminal region.

Next, we quantified the distribution pattern of tagged Bassoon termini at certain distances relative to the TGN. To this end, we defined two distance categories, i.e., 0-100 nm and 101nm—1µm from TGN38 signals. We excluded Bassoon signals farther than 1µm from TGN38 signals, assuming that these signals were not associated with the TGN. The intramolecular N-terminal tags of both double- and single-tagged Bassoon predominantly occupied the 0-100nm distance range. Fraction sizes were 0.67 ±0.1 SD for RFP-Bsn-GFP and 0.84 ±0.04 SD for RFP-Bsn. A smaller fraction of each construct occupied the 101nm—1µm distance range. Here, fraction sizes were 0.33 ±0.1 SD for RFP-Bsn-GFP and 0.15 ±0.04 SD RFP-Bsn (Fig. 4U). In contrast to the intramolecular N-terminal tags, the C-terminal tags of all constructs were more evenly distributed between the two categories, with a tendency towards localization in the 101nm-1µm range: a slightly larger fraction of RFP-Bsn-GFP (0.63±0.1 SD) and Bsn-RFP (0.55±0.03 SD) resided within 101nm—1µm, and the remaining smaller fraction of signals, for double- (0.37±0.1 SD) and single-tagged (0.45 ±0.03 SD) Bassoon constructs, were present within the 0-100nm distance range (Fig. 4U). Thus, all constructs were distributed in such a way that the intramolecular N-terminal tag had a greater likelihood for detection within 100nm of TGN38 than the C-terminal tag. Overall, both the percentage of colocalization and the distance distribution indicate an orientation of recombinant Bassoon at the TGN where the N-termini are arranged closer to the TGN than their C-termini.

Is this arrangement of Bassoon specific for the localization of Bassoon relative to one particular Golgi-protein, i.e., TGN38, or does it reflect the orientation of Bassoon relative to the TGN-compartment in general? To assess this, we imaged RFP-Bsn, Bsn-RFP and RFP-Bsn-EGFP in neurons immunostained for another TGN marker, i.e., Syntaxin6 (Syn6). As seen before with TGN38, the percentage of colocalization with Syn6 was higher for the intramolecular N-terminal tags of RFP-Bsn-EGFP (71.8%) and RFP-Bsn (75.8%) than for the C-terminal tags of RFP-Bsn-EGFP (23.8%) and Bsn-RFP (43.7%; Fig. 5A-Q). The distance analysis also revealed a similar pattern compared to what was seen for TGN38: a significantly larger fraction of intramolecular N-terminal than C-terminal Bassoon signals of single- (0.8±0.03 SD) and double-tagged (0.71±0.02 SD) Bassoon constructs were distributed within the 0-100nm distance range (Fig. 5R). Again, the C-terminal tags of both constructs were more evenly distributed between the two distance categories, and again with a tendency towards localization in the 101nm-1µm range, i.e., a slight majority of C-terminal Bassoon signals of the single- (0.62±0.03 SD) and double-tagged (0.65±0.04 SD) constructs were present in the 101nm—1*μ*m range from the nearest Syn6 signal (Fig. 5R). These results show that Bassoon molecules possess similar colocalization and distance distribution relative to both Syn6 and TGN38 markers of the TGN. This suggests that Bassoon molecules are similarly oriented all over the TGN lamellae, with the N-terminus more closely associated with the TGN membrane than the C-terminus.

### 4.4. Which domains of Bassoon are involved in orienting the molecule?

Next, we aimed at characterizing the features of Bassoon required for bringing the N-terminal region of Bassoon into close apposition to the TGN. N-myristoylation is an obvious possibility for anchoring a protein to membranes. Therefore, we generated a point mutated version of the dually tagged Bassoon construct, called G2A-Bsn, where we replaced glycine in position 2 with alanine. Dual color STED images of TGN38 and the intramolecular N-terminal tag or the C-terminal tag of G2A-Bsn revealed the previously seen pattern of high colocalization of the intramolecular N-terminal tag (74.5%) and low colocalization (28.2%) of the C-terminal tag with TGN38 (Fig. 6A-H, Q). Signals from the intramolecular N-terminal tag were predominantly (fraction size 0.75 ± 0.02 SD) located within 0 - 100nm from the TGN38, while the majority of signals from the C-terminal tag (fraction size 0.75 ± 0.02 SD) were distributed within 101nm- 1µm from the TGN (Fig. 6R). A functional N-myristoylation site is thus not necessary for the orientation of recombinant Bassoon at the TGN. These results do not exclude the possibility that the G2A-mutant recombinant Bassoon dimerizes with endogenous Bassoon, and that the intact endogenous Bassoon somehow helps orienting the N-terminus of the mutant protein. However, when we expressed G2A-Bsn in cultures obtained from mice lacking Bassoon the intramolecular N-terminal tag of G2A-Bsn was still oriented towards the TGN, further corroborating that a functional N-myristoylation site is not necessary for the orientation of Bassoon at the TGN (Suppl. Figure 2).

Since an intact N-myristoylation consensus site was not required for the orientation of Bassoon, we wondered if any other sequences upstream of the intramolecular N-terminal tag were required for the orientation of Bassoon. To test this, we expressed the previously generated GFP-Bsn95-3938 construct aka GFP-95-Bsn. In this construct, the N-terminal 94 amino acids are replaced by EGFP, but it accumulates at the Golgi-apparatus and at synapses (Dresbach et al., 2003; Dresbach et al., 2006). The EGFP tag of this construct, detected by our GFP-nanobody, displayed remarkably low colocalization with TGN38 at the nanoscopical level (Fig. 6 I-L). Its colocalization with TGN38 was 36.2% and the majority of these signals (0.60±0.04 SD) were located farther than 100 nm from the nearest TGN38 signals (Fig. 6Q,R), suggesting that sequences in the first 94 amino acids contribute to orienting the N-terminal area of Bassoon towards the TGN.

A central region of Bassoon, comprising amino acids 2088-2563 and termed Bsn-GBR (for Golgi-binding region), is required for targeting Bassoon to the Golgi-apparatus. Thus, this region may contribute to bringing parts of Bassoon into close proximity to the TGN. A construct consisting of these amino acids fused to the C-terminus of EGFP indeed targets to the Golgi-apparatus, presumably via dimerization with endogenous Bassoon (Dresbach et al., 2006; Maas et al., 2012). Where is this construct located relative to TGN38 at the nanoscopical level? Our STED analysis revealed a relatively low colocalization with TGN38 (41.5%) and predominant signal allocations (0.67±0.03 SD) in the 101nm—1µm distance range from TGN38 signals (Fig. 6. M-P and Q,R). Thus, this region alone, while harboring Golgi-targeting capacity, cannot account for the close apposition of the N-terminal Bassoon regions to the TGN.

Together, these results indicate that neither of the two regions of Bassoon equipped with known Golgi-targeting sequences, i.e., the N-myristoylation consensus site and amino acids 2088-2563, account for the particular orientation of Bassoon at the TGN. Instead, the first 94 amino acids of Bassoon contribute to orienting the N-terminal region of Golgi-associated Bassoon towards the TGN.

#### Estimating the extension of Bassoon molecules at the TGN

Having analyzed the distribution of N- and C-terminal tags over two distance categories, i.e., within 100 nm of TGN38 signals and between 101 nm and 1 µm from the TGN, we wondered whether we might be able to extract the average distance between a tag and TGN38 from the data sets. Figure 7A shows the distances of signals relative to TGN38 obtained from all Bassoon constructs and tags, displayed as violin plots.

The analysis shows that most signals were located within 220 nm from the TGN, irrespective of the construct and tag. Among these signals, two differences between the constructs were obvious: first, the median of the values for distance from TGN38 was smaller for the intramolecular N-terminal tags, both in the dually tagged construct and in the single tagged construct, compared to all other constructs; second, wider sections of the violin plot, representing a higher probability that data in the population take on a certain value, indicated a clustering of intramolecular N-terminal signals close to TGN38. In contrast, all other construct showed a more uniform distribution of signals across the 0-220 nm range (Fig. 7A). We then analyzed the signals from this 0-220 nm range to extract the average distances for each of the constructs and tags from TGN38 (Fig. 7B). In dually tagged Bassoon, average distances from TGN38 were 69 nm for the intramolecular N-terminal tag and 115 nm for the C-terminal tag. Thus, amino acid 97 of Bassoon, where the intramolecular N-terminal tag is located, and amino acid 3938, where the C-terminal tag is located, are estimated to be 46 nm from each other. In single tagged constructs, the intramolecular N-terminal tag was on average 48 nm away from TGN38, the C-terminal tag was 100 nm away. This yields an estimated distance of 52 nm between the average location of the two tags. The average distances from TGN38 of GFP-Bsn95-3938 (106 nm) and Bsn-GBR (115 nm) show that the tags of these deletion constructs occupy similar locations compared to the C-terminal tags of full-length Bassoon constructs (Fig. 7B).

In summary, both transfection of a dually tagged construct and separate transfections of single tagged constructs lead to similar conclusions: first, the intramolecular N-terminal tag is located closer to TGN38 than the C-terminal tag; second, the average distance from TGN38 is between 48 nm and 69 nm for the intramolecular N-terminal tags; the average distance from TG38 is between 100 nm and 115 nm for the C-terminal tags; third, the distance between the intramolecular N-terminal and the C-terminal tag is estimated to be between 46 nm and 52 nm. Overall, this indicates an orderly arrangement of Bassoon molecules with the N-terminus facing the TGN membrane (Fig. 7C).

## 5. Discussion

Bassoon is a presynaptic scaffold protein predicted to be up to 80 nm long (Gundelfinger et al., 2016). At active zones, Bassoon appears to have an extended conformation with its C-terminus facing the plasma membrane (Dani et al., 2010; Limbach et al., 2011). At synapses, Bassoon may thus represent one of the filamentous structures or dense projections characteristic of active zones (Phillips et al., 2001; Dresbach et al., 2001). Here, we find that recombinant Bassoon expressed in cultured hippocampal neurons, has an extended conformation at the Golgi-apparatus. At this subcellular site, the N-terminus of Bassoon faces the TGN membrane. The fact that Bassoon is an extended protein already at this early stage of its trafficking path supports the notion that primordial active zone scaffolds assemble at the Golgi-apparatus (Dresbach et al., 2006). Its orientation relative to membranes - i.e., with the N-terminus facing the TGN membrane, and the C-terminus facing the active zone plasma membrane – adds new insights and raises questions regarding the topology of its trafficking from the Golgi apparatus to active zones.

### 5.1. Design of new Bassoon constructs

We performed this study using a new generation of full-length Bassoon constructs. These constructs were designed to have several advantages. First, the C-terminal tag of these constructs does not produce diffusely distributed fluorescence anymore. This allows for better detection of punctate signals because these are not hidden in a cloud of homogeneous cytoplasmic background fluorescence anymore. We do not know what caused this background in previous constructs (Dresbach et al., 2003). Proteolytic cleavage of the C-terminal tag is a possible cause. But here we found that there is diffusely distributed green fluorescence when the GFP sequence is attached out of frame to the 3’ end of full-length Bassoon, suggesting that a cryptic translation initiation sequence may be present in the 3’ area of the classical construct. By removing the start codon of GFP we eliminated this possibility. Second, to reduce the probability of artificial aggregation we used monomeric versions of fluorescent proteins, i.e., mRFP and the A207K variant of EGFP. Third, all new constructs have a free N-terminus, thus leaving the N-myristoylation consensus site of Bassoon (Dresbach et al., 2003) intact.

To introduce a tag close to the N-terminus, but outside the N-myristoylation consensus sequence, we placed mRFP or mGFP downstream of amino acid 97 in the new Bassoon constructs. In particular, the intramolecular tag is preceded by amino acids 1-97 of Bassoon and followed by amino acids 95-3938 of Bassoon. Originally, this location was chosen because it contains a conveniently located HindIII cloning site in rat Bassoon cDNA. Later it became clear that the first 94 amino acids appear to be a structurally compact unit without persistent folding (Gundelfinger et al., 2016). Glycine and proline constitute 48 percent of the first 94 amino acids of rat Bassoon (25 glycine residues and 20 proline residues), while this percentage steeply drops to 18 percent in amino acids 95-197. Because of the location of the intramolecular tag, the new full-length constructs are expected to combine the properties of two previously characterized constructs: Amino acids 95-3938 preceded by EGFP are correctly targeted to the Golgi-apparatus and to synapses (Dresbach et al., 2003; Bresler et al., 2004; Dresbach et al., 2006; Tsuriel et al., 2009), and amino acids 1-97 followed by EGFP retain targeting capacity for the Golgi-apparatus through their N-myristoylation site (Dresbach et al., 2003; Dresbach et al., 2006). To avoid problems sometimes associated with co-expression, one of our new constructs is dually tagged, with an N-terminal intramolecular mRFP and C-terminal mGFP.

### 5.2. Validating the constructs

All constructs accumulated at the Golgi-apparatus and at presynaptic terminals, as expected (Dresbach et al., 2003; Dresbach et al., 2006). Moreover, their resistance to extraction with Triton X100 indicates their proper incorporation into the CAZ network at synapses (Dresbach et al., 2003). Colocalization levels with the synapse markers Piccolo and Synaptophysin was between 62% and 85%, and is likley underestimated due to rigorous thresholding. The remaining, non-synaptic puncta are probably mobile transport units. This is expected for Bassoon, which traffics on mobile units (Shapira et al., 2003, Bresler et al. 2004) and undergoes activity-induced synapse recruitment to synapses from kinesin-1 – motor dependent axonal carriers (Cai et al., 2007). In addition, we verified by conventional and by STED microscopy that the dually tagged construct was enriched at synapses based on co-localization, in confocal images, with the postsynaptic markers PSD95 and Shank-2/ProSAP1. STED microscopy revealed that this construct was oriented with its C-terminus towards the active zone, as previously shown for endogenous Bassoon (Dani et al., 2010). Thus, we conclude that the novel constructs undergo proper subcellular targeting, that the dually tagged construct adopts the predicted orientation at synapses, and that the two tags can be spatially resolved at subcellular sites where Bassoon has an extended conformation.

Our use of recombinant protein bears the inherent caveat of overexpression. However, recombinant Bassoon has been used widely by us and others to monitor synapse assembly and turnover (Shapira et al., 2003; Cai et al., 2007; Lee et al., 2008; Tsuriel et al., 2009; Matz et al., 2010). In addition, our recombinant proteins accumulated at the expected subcellular sites, and the dually tagged construct showed the expected orientation at the synapse. The design of the constructs also provides several advantages. Under ideal expression conditions the location of the N-terminal intramolecular tag (downstream of amino acid 94) and the C-terminal tag (downstream of amino acid 3938) should report the maximal extension of Bassoon more accurately compared to the routinely used antibodies, whose epitopes are located between amino acids 756 and 1001 and between amino acids 3908 and 3938 (e.g., Dani et al., 2010). Indeed, we observed a distance of 46—52 nm between the tags, compared to 30 nm distance calculated for the two epitopes in Dani et al., 2010. This is consistent with the tags being farther apart within recombinant Bassoon than the two epitopes are in endogenous Bassoon. We emphasize, however, that Dani et al., 2010 performed an extensive 3D analysis of a large number of synapses in brain sections, while we only performed a proof-of-principle analysis for our construct, analysing a set of synapses that appeared to be visible as side views in our cultured neurons.

Using nanobodies to detect the tags provides an additional advantage: because of their small size and direct coupling to fluorophores, nanobodies bring the fluorescent dye closer to the epitope compared to indirect immunofluorescence using primary and secondary antibodies. The subclusters of recombinant Bassoon we detected with nanobodies are consistent with this increased spatial resolution. Maybe these nanobodies detect Bassoon arranged in the presynaptic particle web (Phillips et al., 2001). Overall, we conclude that our analysis of recombinant Bassoon at synapses yields results consistent with previous observations and shows that our approach provides at least similar spatial resolution as previous nanoscopical approaches. Based on these assumptions we conducted our detailed analysis of recombinant Bassoon at the Golgi-apparatus.

### 5.3. Trans Golgi versus trans-Golgi network

STED microscopy revealed that recombinant Bassoon colocalizes with two TGN markers, i.e., TGN38 and Syntaxin-6, but not with a TG sub-compartment marker. This supports and extends previous data showing that both endogenous and recombinant Bassoon colocalize with TGN38 and Syntaxin 6 upon confocal analysis (Dresbach et al., 2006; Maas et al., 2012). This further corroborates that the recombinant protein faithfully represents the localization of endogenous Bassoon. In addition, it narrows down the exact localization of recombinant Bassoon by showing that it is more closely associated with the TGN than with the sub-compartment. This observation is particularly insightful because the recombinant sub-compartment marker we have used here was shown by electron microscopy to selectively label the most “trans” located lamellae of the Golgi stack, i.e., the one immediately preceding the TGN (Wittenmayer, 2014).

### 5.4. Localization and orientation of full-length Bassoon at the trans-Golgi network

A key finding of our study is that the N-terminal, intramolecular tag of Bassoon was located closer to the TGN than the C-terminal tag. This was true both for the dually tagged Bassoon and for the two constructs that carried single mRFP tags. In addition, it was true both when we used TGN38 and when we used Syntaxin-6 as TGN markers. Finally, both colocalization analysis and the distance distribution of the Bassoon constructs indicated this. Thus, recombinant Bassoon is an extended protein located at the TGN, with the N-terminal area closer to the TGN than the C-terminus, and the majority of Bassoon molecules have this orientation.

The distance distribution also revealed that the N-terminal, intramolecular, tag had a higher likelihood of being located within 100 nm nanometers from the TGN than being located inside the 100 nm – to 1 µm range, whereas the C-terminal tag appeared to be distributed more evenly between the two distance regimes. On average, the two tags were located 46 nm away from each other in the dually tagged construct, and 52 nm away from each other when the two single tag constructs were compared. A distance of approximately 50 nm between the intramolecular tag and the C-terminal tag is well within the range of the 80 nm maximal extension of Bassoon predicted by Gundelfinger et al. (2016). It is also consistent with the two tags being farther apart within the primary structure of Bassoon than the two antibody epitopes used in Dani et al. (2010), who estimated a distance of 30 nm between those epitopes based on STORM data. Taken together, our data indicate that recombinant Bassoon is an extended protein located at the TGN, with its N-terminal area oriented towards the TGN membrane and the C-terminus farther away. In addition, they suggest an average distance of 46-52 nm between the intramolecular tag located downstream of amino acid 97 and the C-terminal tag located downstream of amino acid 3938 of rat Bassoon.

### 5.5 Localization and orientation of deletion constructs

Bassoon includes at least two regions with binding capacity for the Golgi-apparatus, including the myristoylated N-terminus and the region spanning amino acids 2088-2563 called the Golgi-binding region (GBR). Using conventional epifluorescence microscopy, we had previously detected constructs encoding either one of these regions at the TGN (Dresbach et al., 2003; Dresbach et al., 2005). The results obtained from analyzing mutated constructs of Bassoon provided some novel insights and raised new questions.

First, the myristoylation-deficient G2A point mutant was still oriented with its N-terminus towards the TGN, even in Bassoon knockout cultures. Thus, insertion of myristic acid into the lipid bilayer is not required for the orientation of the N-terminal region of Bassoon towards the TGN. Through its second coiled-coil domain the G2A-mutant full-length protein may still bind to endogenous Piccolo (Maas et al., 2012). But it is unlikely that this helps orienting the N-terminus of Bassoon towards the TGN, because another construct, GFP-Bsn95-3938, that is also predicted to bind to endogenous Bassoon and Piccolo, showed an aberrant orientation. Thus, N-myristoylation appears to be dispensable for the orientation of Bassoon.

Second, as mentioned above, GFP-Bsn95-3938 was less closely associated with the TGN than the intramolecular tag in the full-length Bassoon constructs. This shows that the N-terminal 97 amino acids of Bassoon are essential for orienting the N-terminus of Bassoon towards the TGN. It is likely that the unusually high percentage of glycine and proline characteristic of this region of Bassoon contributes to this, but the mechanisms and putative binding partners providing this orientation have yet to be discovered. Surprisingly, the tag in GFP-Bsn95-3938 was located unexpectedly far away from the TGN and was rather distributed like the C-terminal tag of the full-length construct. We do not know if some extensive bending of the N-terminal end of this construct towards its C-terminus or a completely aberrant localization account for this.

Third, Bsn-GBR showed a similar localization. This construct includes the second coiled-coil domain of Bassoon, located between amino acids 2246 and 2366 of rat Bassoon (tom Dieck et al., 1998). Bsn-GBR harbors binding capacity for an unknown target site on the Golgi-apparatus and, in addition, dimerizes or oligomerizes with endogenous Bassoon through the CC2 domain (Dresbach et al., 2006; Maas et al., 2012). Therefore, we expected that this construct might bind to the TGN or to a central region of endogenous Bassoon or both. In particular, we aimed to find out if Bsn-GBR locates to the same TGN site that the N-terminus of Bassoon locates to. However, this construct was located farther away from the TGN than the intramolecular tag of Bassoon and had a similarly widespread distribution like the C-terminal tag of full-length Bassoon.

What may account for this localization? Bassoon and Piccolo are required for the biogenesis of Golgi-derived transport vesicles, called gPTVs, that also contain the scaffold protein ELKS2 (Maas et al., 2012). Bassoon binds to ELKS2 through its CC3 regions, to the Golgi apparatus through at least two regions, and to CTBP1, a protein involved in the fission of vesicles budding from the Golgi-apparatus. Thus, Bassoon is endowed with binding capacity for proteins that together could, theoretically, mediate the generation and fission of gPTVs. Overexpressing Bsn-GBR causes the accumulation of Bassoon, Piccolo and ELKS2 at the Golgi-apparatus. Bsn-GBR prevents forward trafficking of gPTVs either by preventing binding of endogenous Bassoon to the Golgi-apparatus, or by impairing oligomerization of Bassoon and possibly Piccolo (Maas et al., 2012). Hence, a complex situation may arise where endogenous proteins gPTVs accumulate and where, in addition, endogenous proteins may be misplaced, when Bsn-GBR is overexpressed. Therefore, the relatively widespread distribution of Bsn-GBR at the nanoscopical level may reflect this construct binding to its target sites in a condition of reduced gPTV exit from the Golgi.

Overall, our results show, that at the nanoscopical level the Bsn-GBR does not bind to the same TGN-region as the N-terminus of Bassoon. Thus, the two Golgi-binding regions of Bassoon seem to associate with distinct sites at the Golgi-apparatus, and the N-terminal 95 amino acids fulfil a special role in orienting the N-terminus towards the TGN.

### 5.6. Perspective: towards a topological scenario

At active zones, the C-terminus of Bassoon is located closer to the plasmamembrane than the N-terminus (Dani et al., 2010; our study). Assuming that Bassoon travels to active zones on Golgi-derived Piccolo-Bassoon transport vesicles (gPTVs; Zhai et al., 2001; Maas et al., 2012) one might predict that the C-terminus of Bassoon is attached to Golgi-membranes and subsequently to the gPTV membrane; deposition of Bassoon at synapses, perhaps by exocytotic fusion of the transport vesicle with the presynaptic plasmamembrane, would then directly place the C-terminus close to the active zone membrane. This scenario is “simple” because it involves no topological rearrangements, i.e. the C-terminus of Bassoon is attached to equivalent membranes all along the trafficking route.

Here, we find that the N-terminus of Bassoon is oriented towards the TGN membrane, while the C-terminus is located farther away from it. In the context of the gPTV model (Zhai et al., 2001; Maas et al., 2012), this is consistent with the following scenario, which also involves no topological rearrangements: at the Golgi-apparatus, the N-terminus of Bassoon may be attached to TGN-associated synaptic vesicle precursor membranes, while the C-terminus may become attached – simultaneously or later – to gPTVs. In this way, Bassoon would travel out of the soma on gPTVs and at the same time carry along synaptic vesicle precursors via its N-terminal region. This speculative scenario is consistent with several observations. First, Bassoon constructs lacking N-terminal areas appear at synapses as small spots the size of Piccolo immunosignals, suggesting that these constructs incorporate into the active zone cytomatrix immediately adjacent to the plasmamembrane; in contrast, a construct comprised of the N-terminal 609 amino acids of Bassoon appears at synapses as larger spots similar to the size of entire synaptic vesicle clusters, suggesting that this N-terminal region of Bassoon may bind to synaptic vesicles (Dresbach et al., 2003). Second, clouds of clear-core vesicles, dense-core vesicles and Bassoon were detected by electron immune-electron microscopy in axons (Tao-Cheng, 2007). It is possible that the dense core vesicles may represent gPTVs while the clear-core vesicles represent synaptic vesicle precursors. Third, a substantial fraction of synaptic vesicles precursors, labelled by recombinant Synaptophysin and called synaptic vesicle transport vesicles (STVs) in this study, co-traffics with active zone precursors, labelled by recombinant Bassoon, as revealed by live imaging studies (Bury and Sabo, 2011; Bury and Sabo, 2015). Recently, a type of Golgi-derived precursor vesicle called PLV, for presynaptic lysosome related vesicles, was identified. PLVs are required for the transport of presynaptic material. They carry, in addition to lysosomal proteins, the synaptic vesicle protein VGlut1 and co-traffic with Bassoon (Vukoja et al., 2018). Thus, at the Golgi-apparatus Bassoon could connect to PTVs via its C-terminus and to synaptic vesicle precursors via its N-terminus, and in this way generate already at the Golgi-apparatus a topological scenario that is later encountered at active zones. Whether this scenario holds true will need to be investigated in future studies.

## Acknowledgements

We are thankful to Prof. Stefan W. Hell, Dr. Dirk Kamin and Dr. Fabian Göttfert for the training and guidance on the STED setups a Max Plank institute of Biophysical Chemistry. We thank Irmgard Weiss for technical assistance. This work was supported by the DFG Research Center for Nanoscale Microscopy and Molecular Physiology of the Brain (CNMPB) to TD.

## References

Acuna, C., Liu, X., and Südhof, T.C. (2016). How to Make an Active Zone: Unexpected Universal Functional Redundancy between RIMs and RIM-BPs. Neuron 91, 792–807.

Bresler, T., Shapira, M., Boeckers, T., Dresbach, T., Futter, M., Garner, C.C., Rosenblum, K., Gundelfinger, E.D., and Ziv, N.E. (2004). Postsynaptic density assembly is fundamentally different from presynaptic active zone assembly. J Neurosci 24, 1507–1520.

Bury, L.A.D., and Sabo, S.L. (2011). Coordinated trafficking of synaptic vesicle and active zone proteins prior to synapse formation. Neural Dev 6, 24.

Bury, L.A.D., and Sabo, S.L. (2016). Building a Terminal: Mechanisms of Presynaptic Development in the CNS. Neuroscientist 22, 372–391.

Cai, Q., Pan, P.-Y., and Sheng, Z.-H. (2007). Syntabulin-kinesin-1 family member 5B-mediated axonal transport contributes to activity-dependent presynaptic assembly. J Neurosci 27, 7284–7296.

Cases-Langhoff, C., Voss, B., Garner, A.M., Appeltauer, U., Takei, K., Kindler, S., Veh, R.W., De Camilli, P., Gundelfinger, E.D., and Garner, C.C. (1996). Piccolo, a novel 420 kDa protein associated with the presynaptic cytomatrix. Eur J Cell Biol 69, 214– 223.

Dani, A., Huang, B., Bergan, J., Dulac, C., and Zhuang, X. (2010). Superresolution imaging of chemical synapses in the brain. Neuron 68, 843–856.

Davydova, D., Marini, C., King, C., Klueva, J., Bischof, F., Romorini, S., Montenegro-Venegas, C., Heine, M., Schneider, R., Schröder, M.S., et al. (2014). Bassoon specifically controls presynaptic P/Q-type Ca(2+) channels via RIM-binding protein. Neuron 82, 181–194.

om Dieck, S., Sanmartí-Vila, L., Langnaese, K., Richter, K., Kindler, S., Soyke, A., Wex, H., Smalla, K.H., Kämpf, U., Fränzer, J.T., et al. (1998). Bassoon, a novel zinc-finger CAG/glutamine-repeat protein selectively localized at the active zone of presynaptic nerve terminals. J Cell Biol 142, 499–509.

Dresbach, T., Qualmann, B., Kessels, M.M., Garner, C.C., and Gundelfinger, E.D. (2001). The presynaptic cytomatrix of brain synapses. Cell Mol Life Sci 58, 94–116.

Dresbach, T., Hempelmann, A., Spilker, C., tom Dieck, S., Altrock, W.D., Zuschratter, W., Garner, C.C., and Gundelfinger, E.D. (2003). Functional regions of the presynaptic cytomatrix protein bassoon: significance for synaptic targeting and cytomatrix anchoring. Mol Cell Neurosci 23, 279–291.

Dresbach, T., Torres, V., Wittenmayer, N., Altrock, W.D., Zamorano, P., Zuschratter, W., Nawrotzki, R., Ziv, N.E., Garner, C.C., and Gundelfinger, E.D. (2006). Assembly of active zone precursor vesicles: obligatory trafficking of presynaptic cytomatrix proteins Bassoon and Piccolo via a trans-Golgi compartment. J Biol Chem 281, 6038–6047.

Fejtova, A., and Gundelfinger, E.D. (2006). Molecular organization and assembly of the presynaptic active zone of neurotransmitter release. Results Probl Cell Differ 43, 49–68.

Fenster, S.D., Chung, W.J., Zhai, R., Cases-Langhoff, C., Voss, B., Garner, A.M., Kaempf, U., Kindler, S., Gundelfinger, E.D., and Garner, C.C. (2000). Piccolo, a presynaptic zinc finger protein structurally related to bassoon. Neuron 25, 203–214.

Garner, C.C., Kindler, S., and Gundelfinger, E.D. (2000). Molecular determinants of presynaptic active zones. Curr Opin Neurobiol 10, 321–327.

Good, M.C., Zalatan, J.G., and Lim, W.A. (2011). Scaffold proteins: hubs for controlling the flow of cellular information. Science 332, 680–686.

Griffiths, G., and Simons, K. (1986). The trans golgi network: sorting at the exit site of the golgi complex. Science 234, 438–443.

Gundelfinger, E.D., Reissner, C., and Garner, C.C. (2015). Role of Bassoon and Piccolo in Assembly and Molecular Organization of the Active Zone. Front Synaptic Neurosci 7, 19.

Hallermann, S., Fejtova, A., Schmidt, H., Weyhersmüller, A., Silver, R.A., Gundelfinger, E.D., and Eilers, J. (2010). Bassoon speeds vesicle reloading at a central excitatory synapse. Neuron 68, 710–723.

Hamers-Casterman, C., Atarhouch, T., Muyldermans, S., Robinson, G., Hamers, C., Songa, E.B., Bendahman, N., and Hamers, R. (1993). Naturally occurring antibodies devoid of light chains. Nature 363, 446–448.

Held, R.G., Liu, C., Ma, K., Ramsey, A.M., Tarr, T.B., De Nola, G., Wang, S.S.H., Wang, J., van den Maagdenberg, A.M.J.M., Schneider, T., et al. (2020). Synapse and Active Zone Assembly in the Absence of Presynaptic Ca2+ Channels and Ca2+ Entry. Neuron 107, 667-683.e9.

Hoffmann-Conaway et al., S., Brockmann, M.M., Schneider, K., Annamneedi, A., Rahman, K.A., Bruns, C., Textori-Taube, K., Trimbuch, T., Smalla, K.-H., Rosenmund, C., Gundelfinger, E.D., Garner, C.C., and Montenegro-Venegas, C. (2020). Parkin contributes to synaptic vesicle autophagy in Bassoon-deficient mice. Elife 9: e56590

Imig, C., Min, S.-W., Krinner, S., Arancillo, M., Rosenmund, C., Südhof, T.C., Rhee, J., Brose, N., and Cooper, B.H. (2014). The morphological and molecular nature of synaptic vesicle priming at presynaptic active zones. Neuron 84, 416–431.

Lee, S.-H., Peng, I.-F., Ng, Y.G., Yanagisawa, M., Bamji, S.X., Elia, L.P., Balsamo, J., Lilien, J., Anastasiadis, P.Z., Ullian, E.M., et al. (2008). Synapses are regulated by the cytoplasmic tyrosine kinase Fer in a pathway mediated by p120catenin, Fer, SHP-2, and beta-catenin. J Cell Biol 183, 893–908.

Limbach, C., Laue, M.M., Wang, X., Hu, B., Thiede, N., Hultqvist, G., and Kilimann, M.W. (2011). Molecular in situ topology of Aczonin/Piccolo and associated proteins at the mammalian neurotransmitter release site. Proc Natl Acad Sci U S A 108, E392–401.

Maas, C., Torres, V.I., Altrock, W.D., Leal-Ortiz, S., Wagh, D., Terry-Lorenzo, R.T., Fejtova, A., Gundelfinger, E.D., Ziv, N.E., and Garner, C.C. (2012). Formation of Golgi-derived active zone precursor vesicles. J Neurosci 32, 11095–11108.

Matz, J., Gilyan, A., Kolar, A., McCarvill, T., and Krueger, S.R. (2010). Rapid structural alterations of the active zone lead to sustained changes in neurotransmitter release. Proc Natl Acad Sci U S A 107, 8836–8841.

Mellman, I., and Simons, K. (1992). The golgi complex: in vitro veritas. Cell 68, 829– 840.

Mendoza Schulz, A., Jing, Z., Sánchez Caro, J.M., Wetzel, F., Dresbach, T., Strenzke, N., Wichmann, C., and Moser, T. (2014). Bassoon-disruption slows vesicle replenishment and induces homeostatic plasticity at a CNS synapse. EMBO J 33, 512– 527.

Montenegro-Venegas, C., Fienko, S., Anni, D., Pina-Fernandez, E., Frischknecht, R., and Fejtova, A. (2021). Bassoon inhibits proteasome activity via interaction with PSMB4. Cell Mol Life Sci 78: 1545–1563.

Okerlund, N.D., Schneider, K., Leal-Ortiz, S., Montenegro-Venegas, C., Kim, S.A., Garner, L.C., Waites, C.L., Gundelfinger, E.D., Reimer, R.J., and Garner, C.C. (2017). Bassoon Controls Presynaptic Autophagy through Atg5. Neuron 93, 897-913.e7.

Phillips, G.R., Huang, J.K., Wang, Y., Tanaka, H., Shapiro, L., Zhang, W., Shan, W.S., Arndt, K., Frank, M., Gordon, R.E., et al. (2001). The presynaptic particle web: ultrastructure, composition, dissolution, and reconstitution. Neuron 32, 63–77.

Sanmartí-Vila, L., tom Dieck, S., Richter, K., Altrock, W., Zhang, L., Volknandt, W., Zimmermann, H., Garner, C.C., Gundelfinger, E.D., and Dresbach, T. (2000). Membrane association of presynaptic cytomatrix protein bassoon. Biochem Biophys Res Commun 275, 43–46.

Schoch, S., and Gundelfinger, E.D. (2006). Molecular organization of the presynaptic active zone. Cell Tissue Res 326, 379–391.

Shapira, M., Zhai, R.G., Dresbach, T., Bresler, T., Torres, V.I., Gundelfinger, E.D., Ziv, N.E., and Garner, C.C. (2003). Unitary assembly of presynaptic active zones from Piccolo-Bassoon transport vesicles. Neuron 38, 237–252.

Südhof, T.C. (2012). The presynaptic active zone. Neuron 75, 11–25.

Tao-Cheng, J.-H. (2007). Ultrastructural localization of active zone and synaptic vesicle proteins in a preassembled multi-vesicle transport aggregate. Neuroscience 150, 575–584.

Tsuriel, S., Geva, R., Zamorano, P., Dresbach, T., Boeckers, T., Gundelfinger, E.D., Garner, C.C., and Ziv, N.E. (2006). Local sharing as a predominant determinant of synaptic matrix molecular dynamics. PLoS Biol 4, e271.

Tsuriel, S., Fisher, A., Wittenmayer, N., Dresbach, T., Garner, C.C., and Ziv, N.E. (2009). Exchange and redistribution dynamics of the cytoskeleton of the active zone molecule bassoon. J Neurosci 29, 351–358.

Vukoja, A., Rey, U., Petzoldt, A.G., Ott, C., Vollweiter, D., Quentin, C., Puchkov, D., Reynolds, E., Lehmann, M., Hohensee, S., et al. (2018). Presynaptic Biogenesis Requires Axonal Transport of Lysosome-Related Vesicles. Neuron 99, 1216-1232.e7.

Waites, C.L., Leal-Ortiz, S.A., Okerlund, N., Dalke, H., Fejtova, A., Altrock, W.D., Gundelfinger, E.D., and Garner, C.C. (2013). Bassoon and Piccolo maintain synapse integrity by regulating protein ubiquitination and degradation. EMBO J 32, 954–969.

Wang, X., Kibschull, M., Laue, M.M., Lichte, B., Petrasch-Parwez, E., and Kilimann, M.W. (1999). Aczonin, a 550-kD putative scaffolding protein of presynaptic active zones, shares homology regions with Rim and Bassoon and binds profilin. J Cell Biol 147, 151–162.

Wildanger, D., Medda, R., Kastrup, L., and Hell, S.W. (2009a). A compact STED microscope providing 3D nanoscale resolution. J Microsc 236, 35–43.

Wildanger, D., Bückers, J., Westphal, V., Hell, S.W., and Kastrup, L. (2009b). A STED microscope aligned by design. Opt Express 17, 16100–16110.

Wittenmayer, N. (2014). Photoconversion of CFP to study neuronal tissue with electron microscopy. Methods Mol Biol 1148, 77–87.

Zhai, R.G., Vardinon-Friedman, H., Cases-Langhoff, C., Becker, B., Gundelfinger, E.D., Ziv, N.E., and Garner, C.C. (2001). Assembling the presynaptic active zone: a characterization of an active one precursor vesicle. Neuron 29, 131–143.

