## Supplemental Figures for "Nanoscopical analysis reveals an orderly arrangement of the presynaptic scaffold protein Bassoon at the Golgi-apparatus"

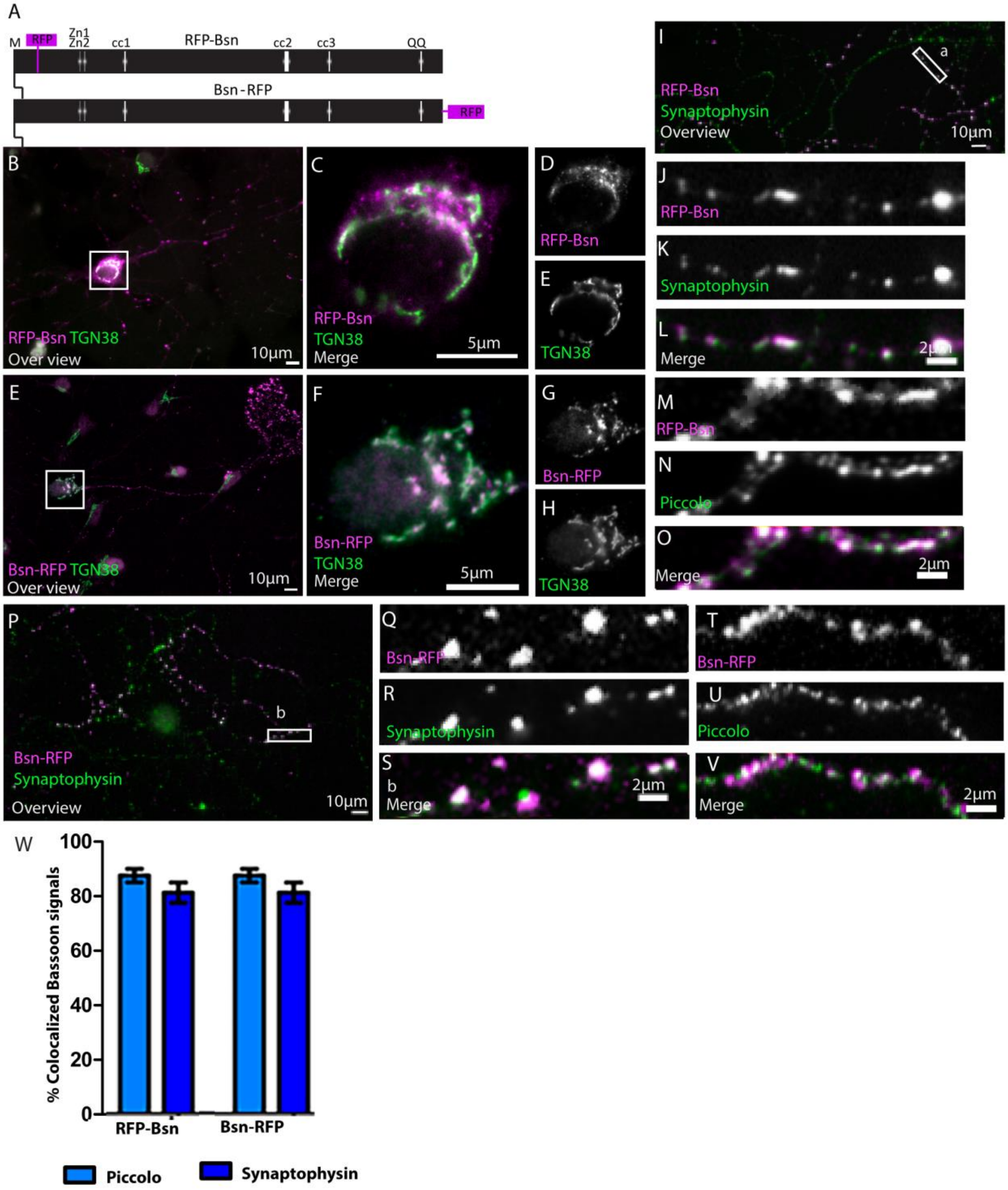


**Supplemental figure 1. Full-length RFP tagged Bassoon constructs localize at the TGN in young neurons, traffick to synaptic sites, and are incorporated into the insoluble AZ scaffold of mature neurons.** Panel **A** is a schematic diagram of full-length Bassoon sequence compared to the sequence of mRFP tagged Bassoon constructs where M stands for N-myristoylation sequence, Zn1 and Zn2 are the two zinc finger domains and cc1, cc2, cc3, are the three predicted coiled-coil regions. Immunostained DIV7 (**B—E& E—H**) and DIV14 (**P—V & I—O**) hippocampal neurons transfected with RFP tagged Bassoon constructs, post a DIV3 lipofectamine transfection, are co-stained with the TGN38 (**B—H**), synaptophysin (**J—L& Q—S**) and piccolo (**M—O & T—V**). Panels **I** and **P** represent 40X over views of the transfections and **a** and **b** represent the zooms of their white square ROIs, respectively. Neurons were fixed in cold methanol prior PFA fixation, to quench the RFP autofluorescence. **W** is the Bassoon colocalization quantification for panels **I—V**; data are represented as mean ± SD, N=5 cells from two separate experiments for each quantification. Scale bars 10 $\mu$ m (**B,** and **E**), 5 $\mu$ m (**C—F**) and 2 $\mu$ m(**L—O** and **S—V**).

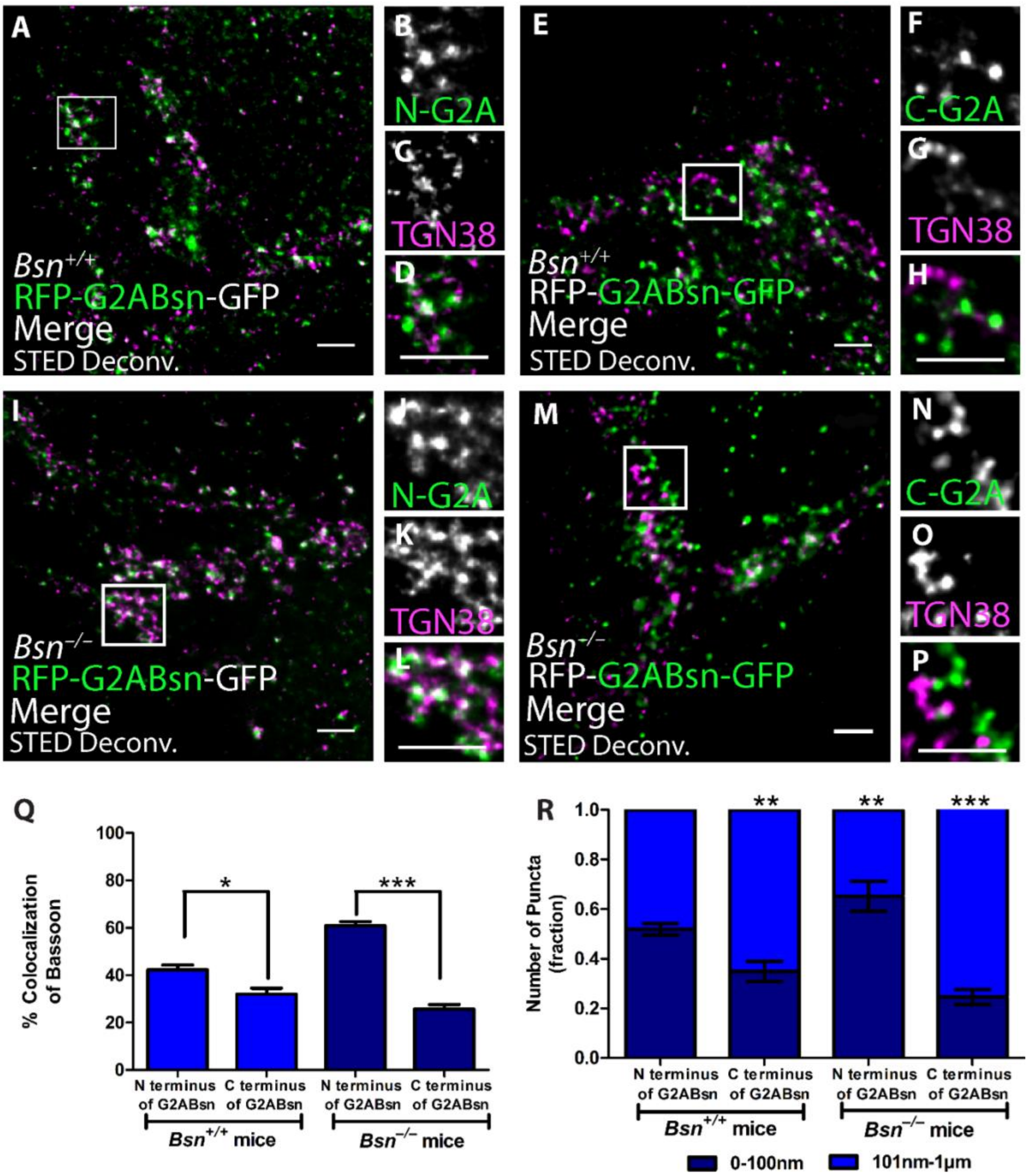


**Supplemental figure 2: Orientation of G2A-mRFP-Bsn-mEGFP myristoyl mutant construct in endogenous Bassoon-free** ***Bsn*$^{-/-}$ knockout mice and their *Bsn*$^{+/+}$ wildtype littermates.**

DIV7 *Bsn*$^{+/+}$ (**A—H**) and *Bsn*$^{-/-}$ (**I—M**) sandwich hippocampal cultures were transfected with G2A-mRFP-Bsn-mEGFP. Two-color STED images with their respective insets are shown for the N– (**A—D** and **I—L**) and C– (**E—H** and **M—P**) termini of the myristoyl mutant construct, respectively. Immunostaining was performed using RFP-nanobody-Atto594 or GFP-nanobody-Atto647 and TGN38 marker. Graph **Q and R** were represented as mean ± SD, N=8 cells from two knockout and two wildtype animals and quantified for amount of colocalization and signal distributions, respectively, $*p \leq 0.05$, $**p \leq 0.01$ and $***p \leq 0.001$. Scale bars 1$\mu$m (**A—P**).
